# Dissecting CD8+ T cell pathology of severe SARS-CoV-2 infection by single-cell epitope mapping

**DOI:** 10.1101/2021.03.03.432690

**Authors:** Felix Schreibing, Monica Hannani, Fabio Ticconi, Eleanor Fewings, James S Nagai, Matthias Begemann, Christoph Kuppe, Ingo Kurth, Jennifer Kranz, Dario Frank, Teresa M Anslinger, Patrick Ziegler, Thomas Kraus, Jürgen Enczmann, Vera Balz, Frank Windhofer, Paul Balfanz, Christian Kurts, Gernot Marx, Nikolaus Marx, Michael Dreher, Rebekka K Schneider, Julio Saez-Rodriguez, Ivan Costa, Rafael Kramann

## Abstract

The current COVID-19 pandemic represents a global challenge. A better understanding of the immune response against SARS-CoV-2 is key to unveil the differences in disease severity and to develop future vaccines targeting novel SARS-CoV-2 variants. Feature barcode technology combined with CITE-seq antibodies and DNA-barcoded peptide-MHC I Dextramer reagents enabled us to identify relevant SARS-CoV-2-derived epitopes and compare epitope-specific CD8^+^ T cell populations between mild and severe COVID-19. We identified a strong CD8^+^ T cell response against an S protein-derived epitope. CD8^+^ effector cells in severe COVID-19 displayed hyperactivation, T cell exhaustion and were missing characteristics of long-lived memory T cells. We identify A*0101 WTAGAAAYY as an immunogenic CD8^+^ T cell epitope with the ability to drive clonal expansion. We provide an in-depth characterization of the CD8^+^ T cell-mediated response to SARS-CoV-2 infection which will be relevant for the development of molecular and targeted therapies and potential adjustments of vaccination strategies.

## Introduction

Since the discovery of Severe Acute Respiratory Syndrome Coronavirus 2 (SARS-CoV-2) in the city of Wuhan, China in December 2019, the pandemic Coronavirus disease 2019 (COVID-19) has posed significant challenges for public health and the global economy at an unprecedented scale^1^. Following the rapid global spread of the highly infectious virus, more than 2 million deaths have been attributed to COVID-19 as of February 2021 (World Health Organization). While most SARS-CoV-2-infected individuals are asymptomatic or display only mild symptoms, some patients develop severe clinical symptoms such as acute respiratory distress syndrome (ARDS), cardiac complications and death^2^. Significant contributions have been made to better understand the infectious disease but the pathogenetic basis for the significant difference in disease severity of COVID-19 is still not well understood.

CD8^+^ cytotoxic T cells play a pivotal role in viral clearance during infection and are activated by the interaction between the T cell receptor (TCR) and virus-derived peptide-antigens presented by major histocompatibility complex (MHC) class I molecules. Following infection, a subset of antigen-specific memory CD8^+^ T cells remains and provides long-lasting protection against secondary viral infection^3^. Efforts have been made to study the role of T cells during SARS-CoV-2 infection and through advances in single-cell RNA sequencing (scRNA-seq) technologies it has become possible to unveil heterogeneity among individual lymphocytes^4^. A thorough characterization of antigen-specific immune cells at a single-cell resolution is crucial for understanding the immunopathology of COVID-19 on a molecular level. While several studies have investigated different characteristics of overall immune cell populations in SARS-CoV-2 infection at a single-cell level^5,6^, only a limited number of studies currently exist that focus on antigen-specific lymphocyte populations, particularly antigen-specific CD8+ T cells^7^. However, studies that combine immune profiling of CD8+ T cells with an examination of epitope-binding properties at a single-cell level are still lacking. A precise understanding of the CD8^+^ T cell response to SARS-CoV-2 epitopes is of particular importance given the current discussions of vaccine efficacy against mutated SARS-CoV-2 variants.

Here, we present scRNA- and TCR-seq profiles of T cells derived from patients with a varying degree of COVID-19 severity with a unique, in-depth characterization of antigen-specific CD8^+^ T cells using DNA-barcoded peptide-MHC I multimers (MHC I Dextramer reagents)^8^ loaded with known SARS-CoV-2-derived peptides. By investigating antigen-specific CD8^+^ T cells we demonstrate that mild cases of COVID-19 display a more functional, terminally differentiated CD8^+^ effector T cell phenotype compared to severe infection. We observe dysregulated interferon (IFN) signaling in severe SARS-CoV-2 infection with a progression towards T cell exhaustion and hyperactivation while antigen-specific CD8^+^ effector cells maintain properties relevant for T cell memory formation in mild cases of COVID-19. Finally, we identify the S protein-derived epitope A*0101 WTAGAAAYY as a potentially important target for CD8^+^ T cell-mediated antiviral response.

## Results

### Identification of immunogenic SARS-CoV-2 epitopes by bulk-sequencing

We first aimed to identify the most immunogenic epitopes from a pool of 38 *in silico*-selected MHC I Dextramer reagents carrying SARS-CoV-2-derived peptides (epitope panel presented in Table S1). This panel also included 4 positive and 4 negative control Dextramer reagents. Patient-specific T cells from the conditions active mild (*n* = 4) and severe (*n* = 4), recovered mild (*n* = 3) and severe (*n* = 11), and healthy controls (*n* = 6) were incubated with the Dextramer reagents followed by subsequent sequencing of Dextramer reagent DNA-barcodes for quantification of epitope-binding T cells and SARS-CoV-2 epitope enrichment (Fig. 1A-B; clinical data presented in Supplementary Table 2).

**Fig. 1.**
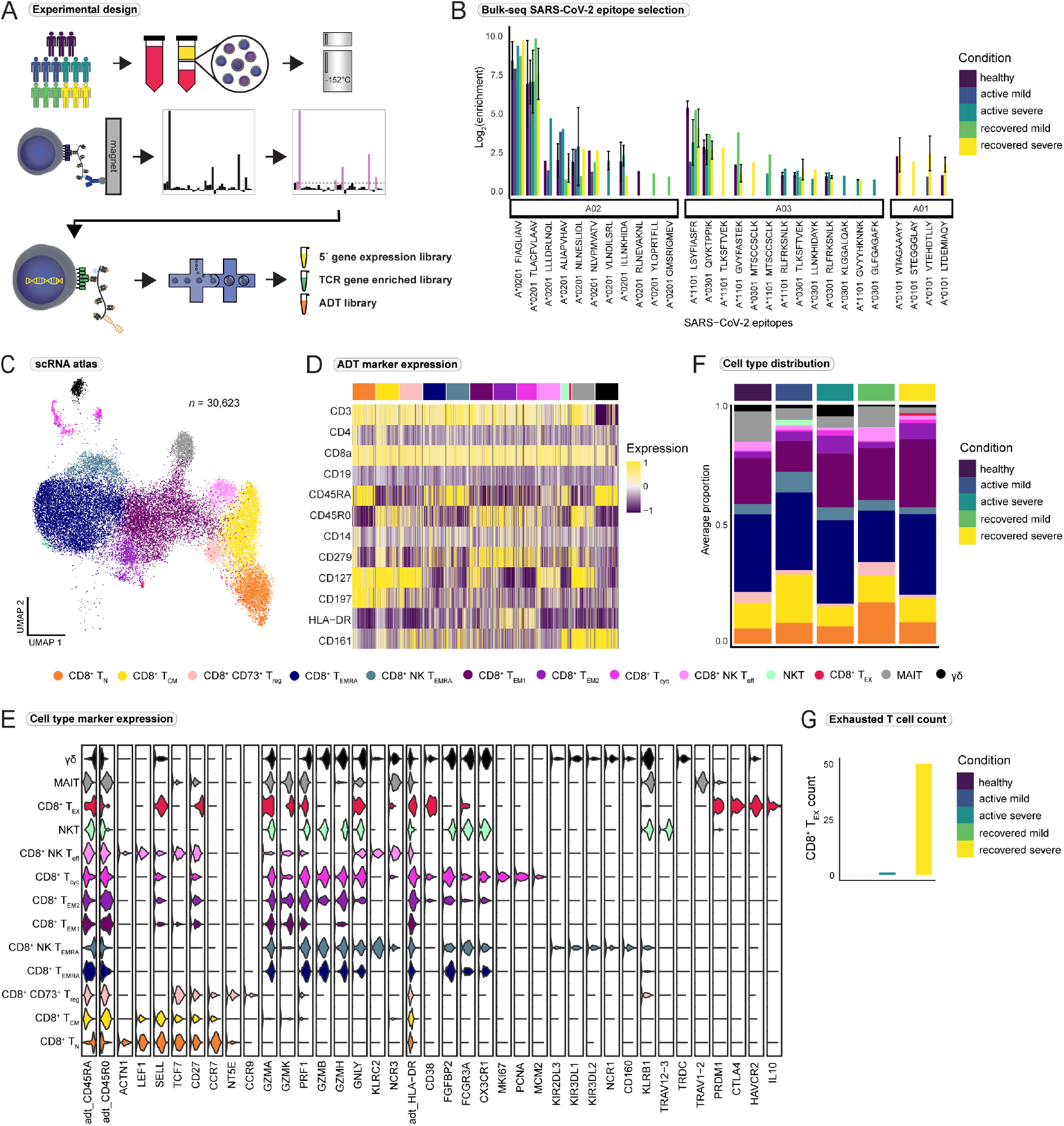
Selection of top SARS-CoV-2 epitopes by bulk sequencing and identification of functional subsets of CD8^+^ T cells. (A) Schematic overview of the study design. (B) Bulk screening enrichment of every peptide-MHC I Dextramer reagent per disease condition plotted as the mean log_2_-fold enrichment. MHC I Dextramer reagents were grouped according to their respective HLA-supertype (see also Supplementary Table 1). (C) Integrated UMAP projection of all 13 CD8^+^ T cell subpopulations (*n* = 30,623). (D) Scaled expression of antibody-derived tag (ADT) markers (CITE-seq) per CD8^+^ T cell subpopulation. (E) Marker gene expression per CD8^+^ T cell subtype. TCR = T cell receptor (see also Supplementary Table 4). (F) Average distribution of CD8^+^ T cell subsets for the healthy (*n* = 2), active mild (*n* = 3), active severe (*n* = 3), recovered mild (*n* = 3), and recovered severe (*n* = 3) condition. Cell type proportions per patient are reported in Supplementary Fig. 2D. (G) Number of exhausted CD8^+^ T cells per condition.

Epitope-binding characteristics were divided into four groups (Supplementary Fig. 1A) based on the HLA profiles of the patients and the Dextramer reagents (Supplementary Fig. 1B-C and Supplementary Table 1 and 3). Patients who displayed the most “specifically enriched” epitope-binding were selected for the subsequent single-cell experiments. Epitopes with an enrichment value of log_2_(5) in at least one patient were considered for further analysis, resulting in 15 top immunogenic SARS-CoV-2 epitopes and 8 controls to form our epitope pool for subsequent single-cell experiments (Supplementary Table 1).

### A population of exhausted CD8^+^ T cells is exclusive for the recovery stage after severe COVID-19

Next, we used the top immunogenic epitopes selected in the bulk screening and performed single-cell immunoprofiling of patient specific CD8^+^ T cells (5’ sequencing, 10x Genomics) by combining scRNA-seq with enrichment of TCR genes and single-cell proteomics (CITE-seq antibodies and MHC I Dextramer reagents carrying SARS-CoV2 epitopes) (Fig. 1A). Patient-specific CD8^+^ T cells were incubated with a panel of immuno-relevant CITE-seq antibodies, a CD8a FACS antibody, and the Dextramer reagents with subsequent FACS sorting (Supplementary Fig. 2A) followed by scRNA-seq.

Unsupervised clustering and subsetting for CD8^+^ T cells captured 30,623 cells and 13 distinct subclusters (Fig. 1C). Functional annotation of T cell subclusters was based on the expression of CD45RA and *CCR7*^9^ together with T cell effector markers^10^ (Fig. 1D-E and Supplementary Table 4). We identified populations of naive CD8^+^ T cells (T_N_), CD8^+^ central memory T cells (T_CM_), and CD73^+^ CD8^+^ regulatory T cells (CD8^+^ CD73^+^ T_reg_) where the latter was annotated based on expression of the regulatory genes *NT5E* (CD73) and *CCR9*^11,12^. A cluster of classic CD8^+^ effector memory T cells re-expressing CD45RA (T_EMRA_) and two populations of CD8^+^ effector memory T cells (T_EM1_ and T_EM2_) were also identified, where T_EM2_ displayed a more activated phenotype based on the expression of *CD38* and *HLA-DR*. A population of cycling CD8^+^ effector cells (T_cyc_) (Supplementary Fig. 2B) also displayed a highly active phenotype. Two distinct clusters strongly expressed natural killer (NK) cell markers *KLRC2* and *NCR3* and were annotated as NK-like CD8^+^ early effector T cell population (NK T_EFF_) and NK-like CD8^+^ effector memory T cell population re-expressing CD45RA (NK T_EMRA_). The NK T_EMRA_ population expressed the killer-cell immunoglobulin-like receptors *KIR2DL3*, *KIR3DL2*, and *KIR3DL1* (Fig. 1E). Of note, we detected a high proportion of CD8^+^ T_EMRA_ cells in one healthy volunteer who informed us of an unknown infection in early January 2020, approximately four months prior to the initiation of this study, which might explain this finding. We therefore decided to remove the volunteer from the study.

We detected three innate-like T cell populations; γδ T cells (γδ), mucosal-associated invariant T cells (MAIT), and atypical NKT cells (NKT) (Supplementary Fig. 2C-D). The NKT cells were characterized by the expression of *TRAV12-3*, *TRBV5-5* and *KLRB1* (CD161) and resembled a population of atypical NKT cells that has been described previously^13^. Interestingly, the group that recovered from severe COVID-19 carried a population of exhausted T cells (T_EX_) (Fig. 1F-G), characterized by the expression of *TIGIT*, *CTLA4*, *CD279* (PD-1) and *HAVCR2* (Tim-3). Exhausted T cells display a strong impairment of effector functions^14^ and are typically seen in chronic infections or tumors, where chronic antigenic stimulation induces exhaustion^15^. This observation indicated that in severe but not in mild COVID-19, T cell exhaustion was induced. Increased number of exhausted T cells during severe SARS-CoV-2 infection is in line with previous findings^16,17^, however conflicting results have recently been reported^7^.

### Impaired interferon response and CD8^+^ T cell differentiation in severe COVID-19

To investigate the mechanisms involved in T cell exhaustion in severe SARS-CoV-2 infection, we focused on the differences between the active disease conditions (active mild vs. active severe infection) and performed differential gene expression and gene set enrichment analysis (GSEA). A significant downregulation of IFN-stimulated genes was found in the active severe condition (Fig. 2A), which was supported by the negative enrichment of biological process (Gene Ontology) gene sets associated with “*response to type I IFN*” in several effector cell types (Fig. 2B).

**Fig. 2.**
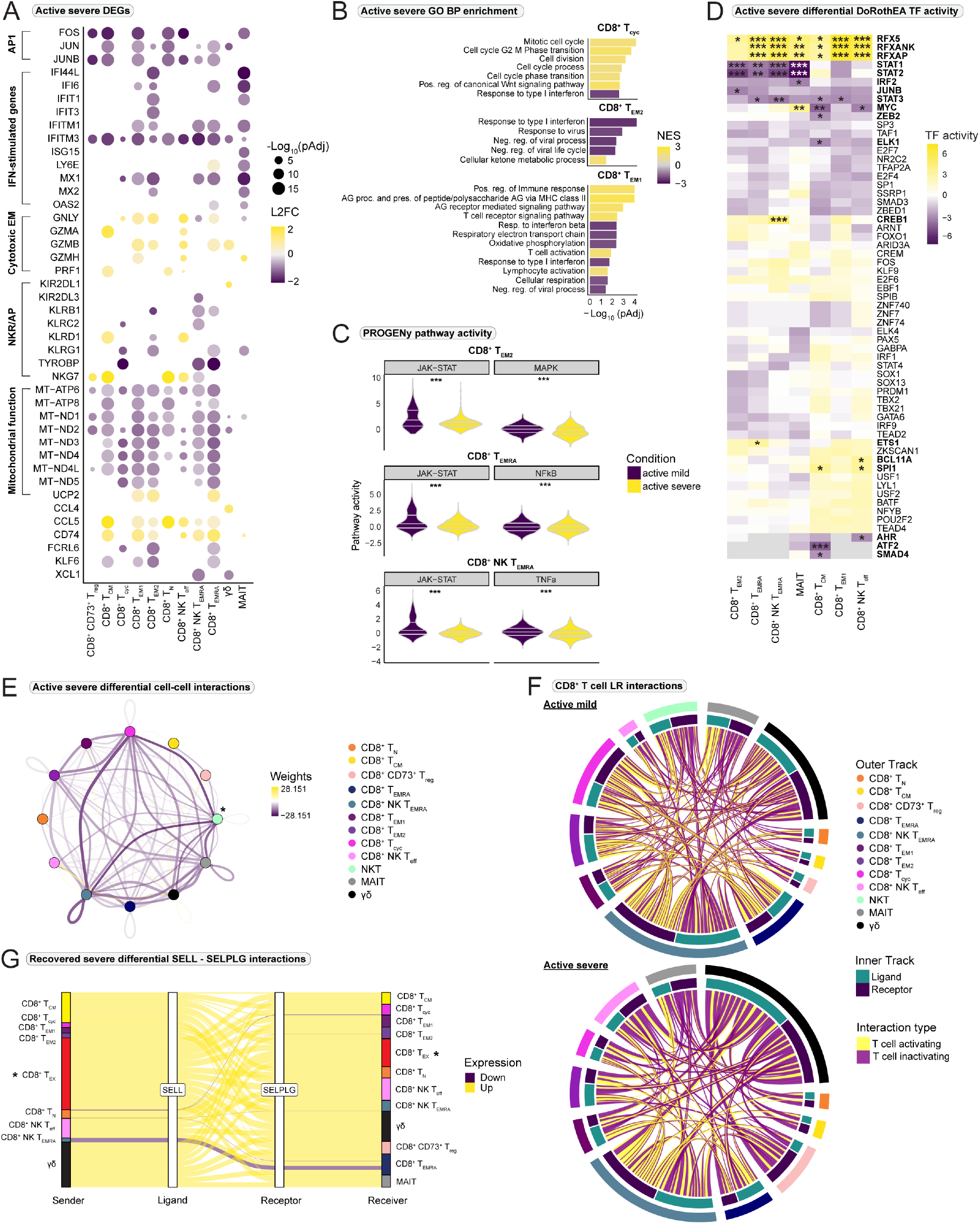
Impaired interferon-response and cell-cell communication in CD8^+^ T cells in active severe COVID-19. (A) Selection of relevant differentially expressed genes (DEG) between active mild and active severe COVID-19. Positive values indicate overexpression in the active severe group. Size and color relate to the adjusted p-value (pAdj) and log_2_ fold change (L2FC), respectively. (B) Gene ontology (GO) Biological process (BP) gene set enrichment analysis. Positive normalized enrichment scores (NES) indicate enrichment in active severe disease. (C) Pathway activity estimated with PROGENy. Selected significant (Wilcoxon rank sum test) pathways are shown (All PROGENy pathways are reported in Supplementary Fig. 3A). (D) Differential transcription factor (TF) activity (DoRothEA) estimated with *msviper*. Positive values indicate increased activity in active severe COVID-19. Significant genes are highlighted by name. (E) Differential cell-cell interactions between active severe and mild COVID-19. Negative values indicate fewer interactions in the active severe group. *NKT cells were only included for the active mild condition. (F) Selected T cell activating (yellow) and inhibiting (purple) ligand-receptor (LR) interactions between CD8^+^ T cell subtypes. Supplementary Fig. 3C is referred to for specific interactions included. (G) Sankey plot of differential interactions of SELL-SELPLG between recovered severe and mild COVID-19. Positive values indicate increased interactions in the recovered severe group. *T_EX_ cells were only present in the recovered severe condition. IFN = interferon, EM = effector molecule, NKR = NK cell receptor, AP = adaptor protein, pos = positive, neg = negative, reg = regulation. *** = p < 0.001, ** = p < 0.01, * = p < 0.05.

Previous studies have demonstrated that JAK-STAT signaling is crucial for the cellular response to IFN-α and −β as activated STAT1/STAT2 heterodimers bind to IFN regulatory factor 9 (IRF9) to form the IFN stimulated gene factor 3 (ISGF3)^18^. ISGF3 translocates to the nucleus to initiate transcription of other IFN regulatory factors (IRFs), which in turn induce the expression of IFN-stimulated genes^19–21^. To investigate whether impaired JAK-STAT signaling could be the reason for the reduced expression of IFN-stimulated genes in severe COVID-19, we estimated the signaling pathway and transcription factor activity. The analysis revealed significantly stronger JAK-STAT pathway activity in differentiated effector cell types from mildly affected individuals (Fig. 2C). This was further supported by a significantly stronger transcription factor activity of STAT1 and STAT2 (Fig. 2D). Among the IFN-stimulated genes, *IFITM3* was significantly downregulated in all CD8^+^ T cell subtypes in active severe COVID-19 when compared to mild (Fig. 2A).

A significant underexpression of the genes *KLRG1* and *FCRL6* was observed in effector cell subtypes in the active severe condition (Fig. 2A), which have been associated with highly differentiated effector phenotypes of CD8^+^ T cells^22,23^. Genes encoding mitochondrial respiratory chain complex I were significantly downregulated in severe disease whereas mitochondrial uncoupling protein 2 (*UCP2*) was overexpressed. Inhibition of complex I has been associated with impaired CD8^+^ T cell effector function as it is important for metabolic reprogramming upon T cell activation^24^, while *UCP2* reduces mitochondrial oxidative phosphorylation and restricts terminal differentiation of CD8^+^ T cells towards short-lived effector cells^25^. While terminally differentiated effector T cells are present in both the active mild and severe condition, these findings indicate impaired effector differentiation and a short-lived nature of CD8^+^ effector T cells in severe COVID-19. In this context, overexpression of *UCP2* could be interpreted as a negative feedback intended to counteract terminal differentiation and a short-lived phenotype. Significantly stronger activity of RFX transcription factors was observed in active severe SARS-CoV-2 infection (Fig. 2D). These transcription factors have been shown to facilitate transcription of MHC class II genes^26^. Expression of MHC class II molecules is a sign of T cell activation^27^ therefore stronger transcription of these genes could be indicative of hyperactivation in severe COVID-19. Finally, several cytolytic effector molecules were upregulated in CD8^+^ effector T cells (Fig. 2B). In summary, these results suggest that CD8^+^ effector T cells in severe COVID-19 display a highly cytotoxic and short-lived phenotype with characteristics of hyperactivation. A full list of differentially expressed genes and an overview of all investigated pathways are provided in Supplementary Table 5 and Supplementary Fig. 3A.

### Altered cell-cell communication promotes exhaustion and impaired effector T cell survival in severe COVID-19

We next asked whether cell-cell communication contributed to an impaired CD8^+^ T cell response in active severe COVID-19. Differences in the global interactions between active mild and severe SARS-CoV-2 infection revealed an overall decrease in cell-cell communication in the severe condition (Fig. 2E and Supplementary Fig. 3B). We focused on a selection of ligand-receptor interactions that have been associated with effector T cell activation or inhibition and displayed a high difference between the active disease conditions (Fig. 2F). These included killer-cell immunoglobulin-like receptors (KIRs) and lectin-like receptors. A large decrease in activating and inhibiting receptor signaling in the T_cyc_ and T_EMRA_ populations was observed for the active severe group. Larger heterogeneity and increased average signaling strength of interactions via several lectin-like receptors (NKG2A, NKG2C, NKG2D, NKG2E, CLEC2B, and KLRB1) towards CD8^+^ effector T populations were observed for the active mild group (Supplementary Fig. 3C). Of note, signaling towards T_cyc_ via CD94/NKG2A was missing in the active severe condition (Supplementary Fig. 3D). The heterodimer CD94/NKG2A is known to inhibit effector functions of CD8^+^ T cells^28,29^ and NKG2A-knockout in influenza- and adenovirus-infected mice has been shown to increase lung pathology^28^. Therefore, lack of NKG2A-mediated inhibition could contribute to immunopathology, hyperactivation of CD8^+^ effector T cells and lung injury in severe COVID-19. Engagement of NKG2A has also been reported to increase survival of virus-specific CD8^+^ effector T cells^29,30^, supporting the finding of short-lived CD8^+^ effector T cells in severe COVID-19. Interestingly, the average strength of MICB-NKG2D signaling towards T_EM2_ and T_EMRA_ cells was stronger in the active severe condition (Supplementary Fig. 3D). As NKG2D is known to mediate costimulatory signals upon TCR ligation^31^, this might contribute to a hyperactivated effector phenotype in severe COVID-19.

We observed stronger predicted interactions via KIRs in active mild COVID-19 towards the NK T_EMRA_ and T_cyc_ populations than in severe COVID-19 (Supplementary Fig. 3C). In contrast, predicted strength of interactions towards γδ T cells via KIR2DL1 and KIR3DL2 was increased in severe COVID-19. Besides their regulatory effect on CD8^+^ T cell activation, several human and murine studies have suggested that KIRs play an important role in long-lived CD8^+^ T cell memory formation^32,33^, possibly by inhibiting activation-induced cell death^32,34^. These results could suggest a more potent memory induction in mild COVID-19.

Increased average signaling strength was found for SELL-SELPLG interactions in most CD8^+^ T cell populations in active severe COVID-19 (Supplementary Fig. 3D). SELPLG has been shown to induce CD8^+^ T cell exhaustion in mice infected with lymphocytic choriomeningitis virus (LCMV). Furthermore, *SELPLG*-knockout was associated with an increase in IL-7Rα-expressing antigen-specific T cells that displayed increased survival^35^. After acute LCMV infection, *SELPLG*-knockout mice displayed increased numbers of epitope-specific CD8^+^ memory T cells^36^. Additionally, we observed IL-10 signaling towards all CD8^+^ T cell subtypes in the population that recovered from severe COVID-19. Increased IL-10 production has been observed in murine models of T cell exhaustion^37^ and could be attributed to a population of severely exhausted CD8^+^ T cells^38^. In line with this, exhausted CD8^+^ T cells in the recovered severe condition displayed strong interactions with other CD8^+^ T cell populations via SELL-SELPLG, thereby contributing to further exhaustion (Fig. 2G).

In summary, our results support an imbalance in CD8^+^ T cell regulation via NK cell receptors in severe COVID-19. This might contribute to a hyperactivated and short-lived effector phenotype. Changes in cell-cell interactions appeared to promote CD8^+^ T cell exhaustion in severe SARS-CoV-2 infection, while exhausted T cells themselves further contributed to the process of exhaustion by interacting with other CD8^+^ T cell populations. Furthermore, the cell-cell interaction analysis suggested a decreased survival and impaired memory formation in favor of exhaustion in severe COVID-19.

### Severe COVID-19 displays shift towards terminal T cell differentiation and exhaustion

To understand potential differences in CD8^+^ T cell differentiation between active mild and severe COVID-19, we performed a pseudotime trajectory analysis on CD8^+^ T cells that would likely originate from naive CD8^+^ T cells (*n* = 27,769). Pseudotime analysis predicted two trajectories originating from the T_N_ population (Fig. 3A). The first trajectory progressed towards the terminally differentiated T_EMRA_ stage and was designated the short-lived effector cell (SLEC) lineage. The second trajectory progressed towards the T_EM2_ and T_EX_ populations and was termed the memory-precursor effector cell (MPEC) lineage (Fig. 3A). We observed a significant shift in cell density for active severe disease towards the late stages of pseudotime in both trajectories compared to mild COVID-19 (Fig. 3B and Supplementary Fig. 4A). This could indicate an imbalanced shift towards terminal T cell effector differentiation and exhaustion in severe SARS-CoV-2 infection.

**Fig. 3.**
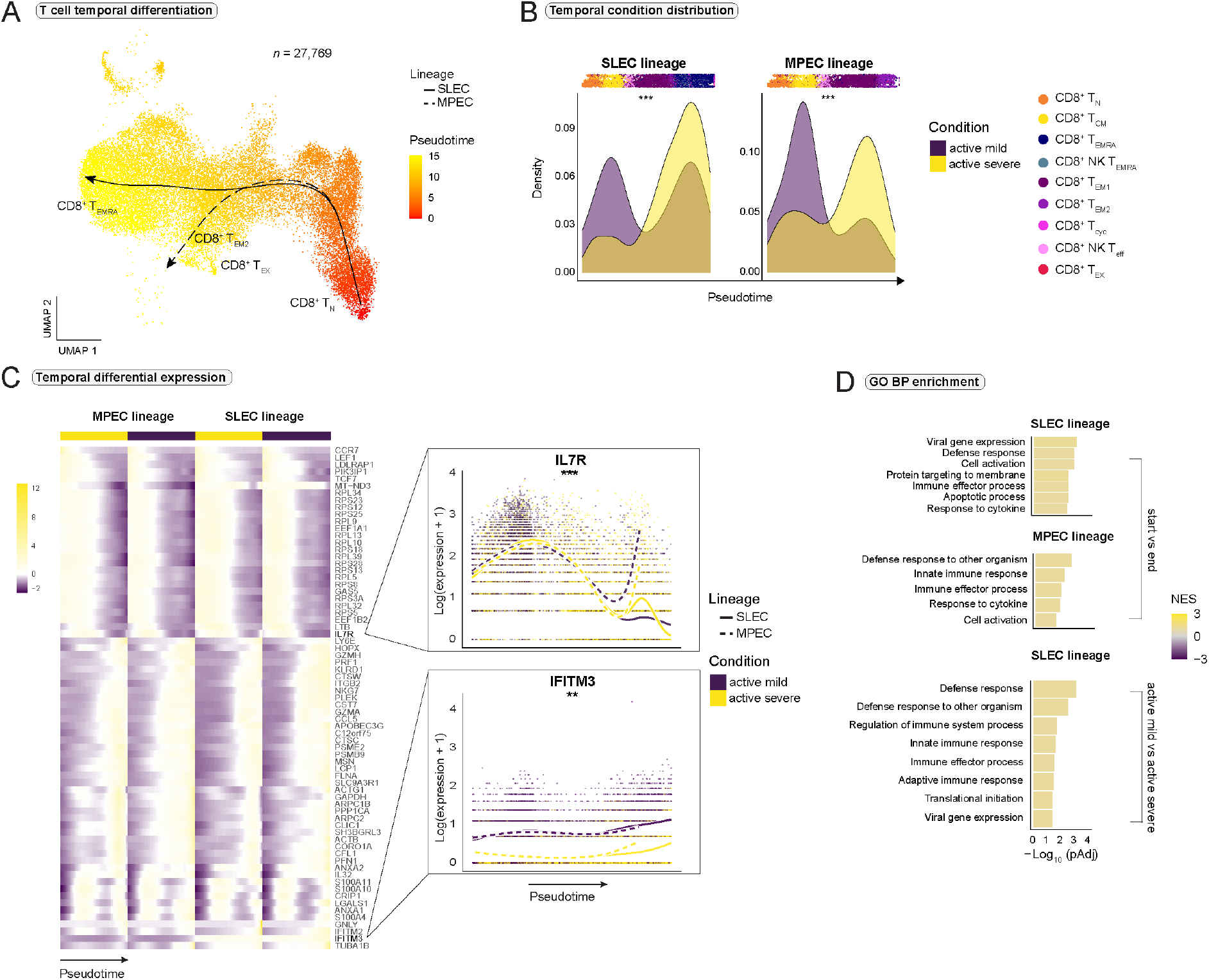
Impaired CD8^+^ T cell differentiation and exhaustion in severe COVID-19. (A) Pseudotimes and estimated trajectories projected onto the integrated UMAP of cell types likely to have their origin in naive CD8^+^ T_N_ cells. (B) Temporal distribution of cell density for the active mild and severe conditions across pseudotime. Shifts in distribution between the conditions for the short-lived effector cells (SLEC) and memory precursor effector cells (MPEC) lineage were tested with the Kolmogorov-Smirnov method (MPEC: D = 0.310, p < 2e-16, SLEC: D = 0.402, p < 2e-16). Supplementary Fig. 4A is referred to for distribution of healthy and recovered conditions. (C) Heatmap depicts differentially expressed genes between the progenitor and differentiated cell populations across pseudotime (start vs end). Smoothed expression of two selected genes is shown with the y-axis on natural logarithmic scale. An extended panel of genes is reported in Supplementary Fig. 4. (D) Significantly enriched biological process (BP) gene sets from the Gene Ontology (GO) database. Gene set enrichment analysis was based on the genes from the start vs end test (upper panel) and condition tests between active mild and severe COVID-19 (lower panel) with *tradeSeq*. Normalized enrichment score = NES. *** = p-value < 0.001, ** = p-value < 0.01.

To dissect differences between the SLEC and MPEC trajectories and between the active mild and severe disease states within the lineages, temporal differential gene expression was conducted (Fig. 3C). Differential gene expression between the progenitor and differentiated cell populations revealed a significant increase in the expression of cytotoxic effector molecules over pseudotime in both lineages, indicative of acquisition of effector functions. The SLEC lineage was significantly enriched for the biological processes “*cell activation*”, “*immune effector process*”, and “*apoptotic process*”, which was indicative of acquisition of a short-lived effector phenotype at the end of this trajectory. Interestingly, genes associated with “*viral gene expression*” were also significantly enriched in the SLEC lineage (Fig. 3D).

To decipher functional differences between the SLEC and MPEC lineages, we performed differential expression analysis in the pseudotime partition where the two lineages bifurcated (Supplementary Table 5). A strong increase in the expression of *KLRC2* was observed from the bifurcation point in the SLEC lineage (Supplementary Fig. 4B). *KLRC2* encodes for the activating NK cell receptor NKG2C and has been shown to be expressed in highly differentiated T_EMRA_ cells^39^, again indicating a terminal effector differentiation in this trajectory. In contrast, we observed a re-expression of *IL7R* from the bifurcation point in the MPEC lineage after an initial decrease in expression (Fig. 3C). The same expression pattern for IL-7Rα expression has been reported in memory precursor cells, evolved from antigen-induced effector cells, that developed into long-lived memory cells^23^. This suggested a memory-precursor profile of the MPEC lineage where a subset of effector cells is able to re-express IL7-Rα and differentiate into long-lived memory cells^40^. However, it should be noted that the T_EX_ population was also present at the end stage of the MPEC lineage. Interestingly, increased IFN-stimulated gene expression was observed in the end stage of the MPEC lineage compared to the SLEC trajectory (Supplementary Fig. 4C), indicating elevated IFN-responsiveness of these cells. Thus, we hypothesised the MPEC lineage comprised a strongly IFN-responsive memory precursor effector cell population that terminated with exhaustion in severe COVID-19.

### Potentially reduced viral protection in severe COVID-19

To elucidate differences in functional CD8^+^ T cell activation between active mild and severe SARS-CoV-2 infection, we compared the active disease conditions within the SLEC and MPEC lineages. Several IFN-stimulated genes displayed increased expression in the active mild condition compared to severe, including the gene *IFITM3* (Fig. 3C). IFITM3 has previously been reported to confer protection against H1N1 influenza A virus^41,42^ and IFITM proteins have been shown to restrict cellular entry of SARS-CoV-1^43^. Therefore, our finding could indicate a potential role for IFITM3 in protection from severe courses of SARS-CoV-2 infection. Several MHC II genes were significantly upregulated in both lineages in active severe COVID-19 compared to mild Elevated MHC II expression in T cells has previously been described as a sign of activation^27^, supporting our finding of CD8^+^ T cell hyperactivation in severe COVID-19. Conversely, several activating NK receptor genes were significantly upregulated in the SLEC and MPEC trajectories in the active mild condition, potentially indicating more effective T cell effector differentiation during mild COVID-19.

### TCR analysis reveals clonal hyperexpansion in severe COVID-19

To investigate the characteristics of clonal expansion of CD8^+^ T cells in the COVID-19 conditions, we analysed the single cell TCR-seq data. We observed an increase in clonal expansion from early to the late stages of CD8^+^ T cell differentiation (Fig. 4A) and found that the differentiated CD8^+^ T cell populations were composed of larger clonotypes (Fig. 4B and Supplementary Fig. 5A). Furthermore, we observed a high proportion of hyperexpanded clones in active severe COVID-19 (Fig. 4B and Supplementary Fig. 5A).

**Fig. 4.**
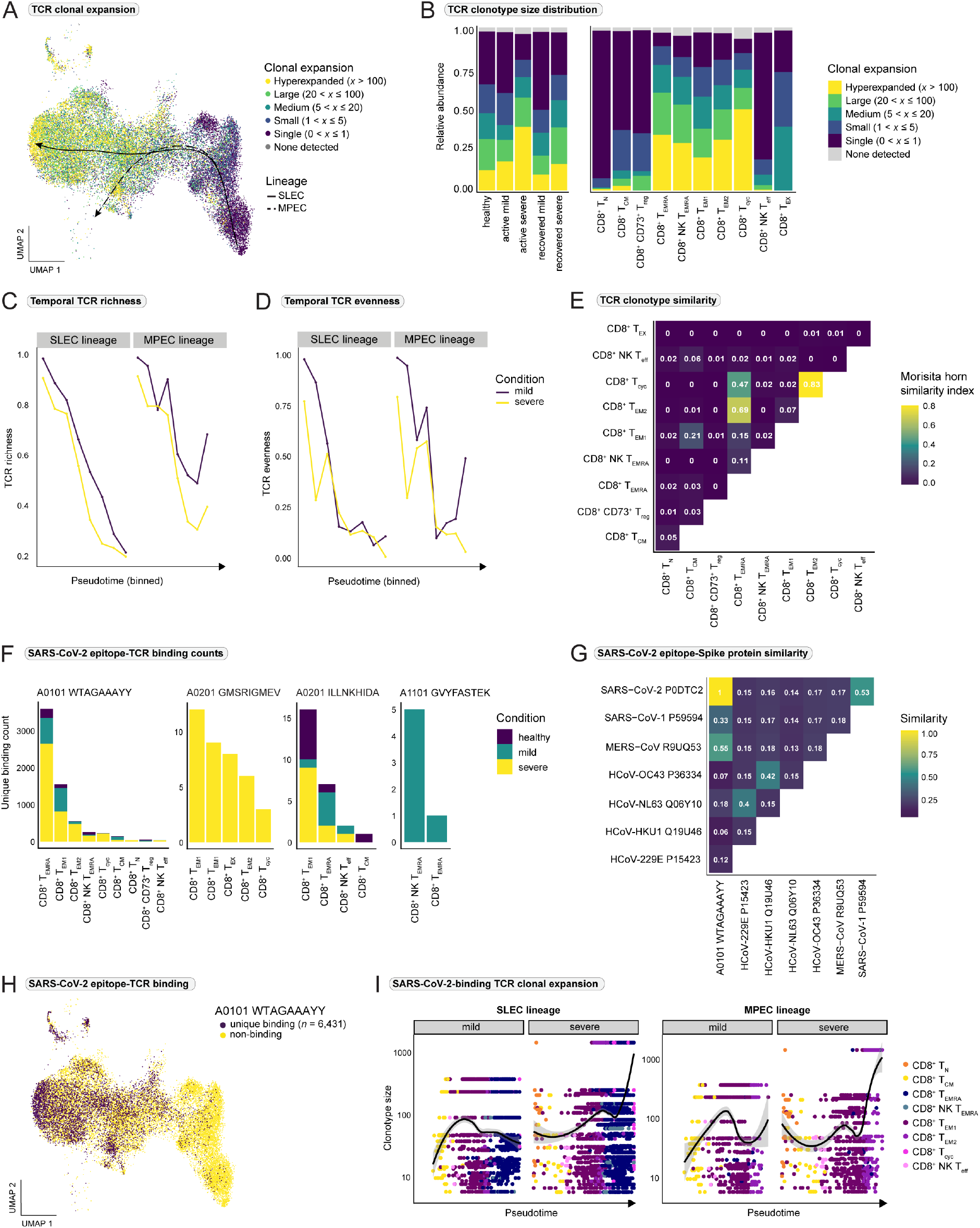
Clonal hyperexpansion is pronounced in severe COVID-19 and CD8^+^ T cells highly respond to viral Spike-protein-derived WTAGAAAYY. (A) T cell receptor (TCR) clonal expansion projected onto the integrated UMAP of cell types, which are likely to have their origin in naive CD8^+^ T_N_ cells. (B) Distribution of clonal expansion within the conditions (left) and within the CD8^+^ T cell populations (right), displayed as relative abundance of clonotype expansion groupings (Supplementary Fig. 5A is referred to for abundance per cell type per condition). (C) TCR richness and (D) evenness across pseudotime for the short-lived effector cell (SLEC) and memory precursor effector cell (MPEC) trajectories for the collapsed mild and severe conditions (active and recovered). (E) TCR clonotype similarity between the CD8^+^ T cell subpopulations estimated with the Morisita horn similarity index. (F) Binding counts for the four uniquely recognized SARS-CoV-2-derived epitopes per CD8^+^ T cell subtype and condition (Supplementary Fig. 5B is referred to for binding counts per patient). (G) Protein similarity between the Spike protein-derived epitope WTAGAAAYY and Spike proteins from other known human coronaviruses. (H) Unique WTAGAAAYY epitope-binding cells (*n* = 6,431) highlighted on the integrated UMAP (Supplementary Fig. 5C highlights the top four WTAGAAAYY epitope-binding clonotypes on the integrated UMAP). (I) Clonal expansion of WTAGAAAYY epitope-binding CD8^+^ T cells across pseudotime per condition and trajectory.

In order to study global changes in diversity of the TCR repertoire during COVID-19, TCR richness and evenness were computed for the collapsed mild and severe conditions (active and recovered) (Fig. 4C-D). We observed a global decrease in TCR diversity over pseudotime for the SLEC and MPEC lineages for both mild and severe COVID-19. However, the severe condition displayed a more accentuated loss of diversity in the TCR repertoire even from early stages of pseudotime, which was in line with the observation of hyperexpansion in this group. To study the similarities between the TCR repertoires of the overall CD8^+^ T cell subpopulations, we computed the overlap in clonotype abundance between the CD8^+^ T cell subtypes. T_EM2_ displayed high similarity to T_cyc_ and the TCR repertoire of T_EMRA_ was similar to both T_EM2_ and T_cyc_ (Fig. 4E). As mainly antigen-induced T cells enter the cell cycle, the observed similarity could indicate that the T_EM2_ and T_EMRA_ populations consisted of clonotypes that received strong proliferative stimuli and could be of importance for the antigen-specific response to COVID-19.

### CD8^+^ T cells highly respond to a SARS-CoV-2 Spike-protein-derived epitope

To unveil SARS-CoV-2-derived epitopes that may drive clonal expansion of CD8^+^ T cells and to investigate differences in epitope-specific CD8^+^ T cells between mild and severe COVID-19, we integrated the scRNA- and TCR-seq data with epitope-binding information from the pool of MHC I Dextramer reagents loaded with known SARS-CoV-2-derived peptides. We identified eight epitopes that were bound by CD8^+^ T cells (Supplementary Table 6) of which four were uniquely recognized by cells that did not bind any other epitope (Fig. 4F). Of these four, two epitopes were derived from the SARS-CoV-2 S protein (*WTAGAAAYY* and *GVYFASTEK*) and the remaining two from the N protein (*GMSRIGMEV* and *ILLNKHIDA*) (Supplementary Fig. 5B). Most of the uniquely epitope-binding cells were specific for WTAGAAAYY, which indicates a relevance of this S protein-derived epitope for CD8^+^ T cell-mediated immunity to SARS-CoV-2. Therefore, we investigated the amino acid sequence similarity between the peptide and S proteins derived from other human coronaviruses (HCoVs). The peptide displayed high similarity to MERS-CoV and SARS-CoV-1 but low similarity to the “common cold” HCoVs (Fig. 4G).

Unique A*0101 WTAGAAAYY epitope-binding was distributed across most subpopulations of MHC class I restricted CD8^+^ T cells (*n* = 6,431) and was highly enriched in the differentiated populations (Fig. 4F and 4H). This finding was supported by increased clonal expansion of WTAGAAAYY epitope-specific CD8^+^ T cells over pseudotime for both the SLEC and MPEC trajectories (Fig. 4I). For severe COVID-19, a strongly hyperexpanded epitope-binding clonotype (*n* = 1,499) was revealed (Supplementary Fig. 5C) and its functional development could be followed from the naive state T_N_ to terminal differentiation (Fig. 4I). TCR-seq analysis revealed that the TCRɑ (TRA) and TCRβ (TRB) complementary determining region 3 (CDR3) sequences of the strongly hyperexpanded clonotype from the severe group (CILYRYDKVIF;CASSLEAGADPVHRQFF) made up approximately 25% of the TRA/B usage in this group (Fig. 5A-B). In contrast, the most frequently used CDR3s in mild COVID-19 represented nearly 5% (CAPLAGSNYQLIW;CASSVTDSSYEQYF) of the TRA/B usage in this group. This further supported the observation of a more accentuated clonal hyperexpansion of CD8^+^ T cells upon A*0101 WTAGAAAYY epitope-recognition in severe COVID-19. MHC I-binding predictions revealed that seven out of 12 patients whose CD8^+^ T cells recognized A*0101 WTAGAAAYY would be able to present this epitope on their endogenous MHC I molecules.

**Fig. 5.**
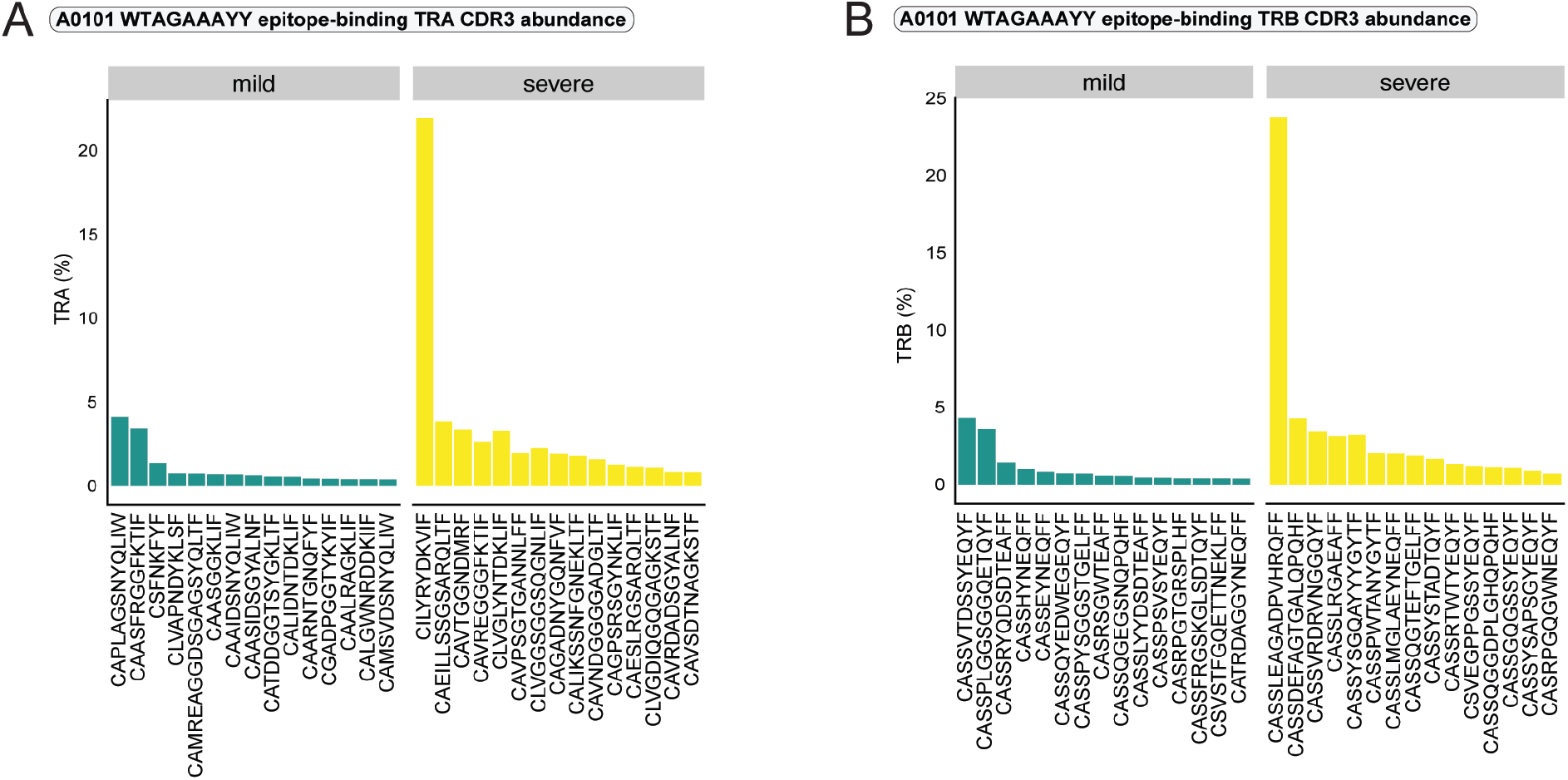
Pronounced clonal hyperexpansion of SARS-CoV-2-specific CD8^+^ effector T cells during severe COVID-19. (A) Percentage of complementary determining region 3 (CDR3) usage of the 15 most abundant clonotypes per collapsed disease condition (active and recovered) for the T cell receptor ɑ chain (TRA) and (B) β chain (TRB).

### Exhaustion and impaired memory formation of epitope-specific CD8^+^ T cells in severe COVID-19

In order to elucidate the differences between the SARS-CoV-2 epitope-specific CD8^+^ T cells in mild and severe COVID-19, we focused on the uniquely WTAGAAAYY epitope-binding T_EM1_ and T_EMRA_ populations. Differential gene expression analysis between mild and severe COVID-19 revealed an upregulation of certain IFN-stimulated genes in the epitope-binding cells of the severe group (Fig. 6A and Supplementary Table 5). Additionally, JAK-STAT pathway activity was significantly stronger in both epitope-binding populations in severe COVID-19 (Fig. 6B and Supplementary Fig. 5D). Enhanced JAK-STAT signaling activity was most likely causative for the stronger expression of IFN-stimulated genes in the epitope-binding T_EM1_ and T_EMRA_ cells in severe COVID-19. These findings were in contrast to the observations from the overall CD8^+^ T cell subsets (Fig. 2A-C). Interestingly, we also observed a significant upregulation of *STAT1* expression and downregulation of *STAT4* expression in WTAGAAAYY epitope-binding T_EMRA_ cells in severe COVID-19 compared to mild. However, *IFITM3* was still significantly downregulated in the antigen-specific T_EM1_ cells in severe disease compared to mild COVID-19, further indicating a special relevance of IFITM3 for protection against severe SARS-CoV-2 infection.

**Fig. 6.**
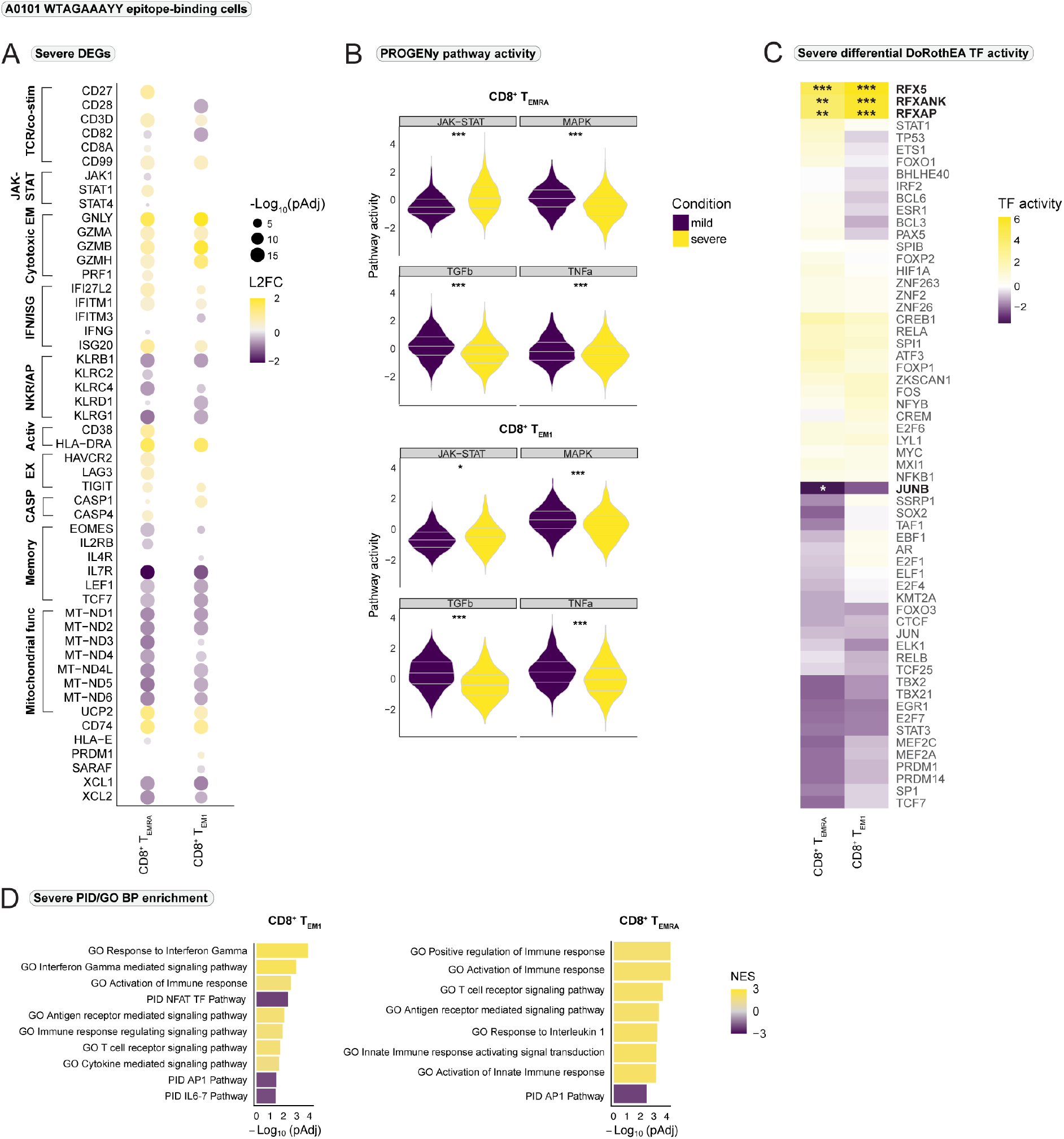
WTAGAAAYY epitope-binding CD8+ T cells display a hyperactivated and less differentiated effector phenotype in severe COVID-19. (A) Differentially expressed genes (DEG) in WTAGAAAYY epitope-binding CD8^+^ T cells between severe and mild COVID-19. Positive values indicate overexpression in the severe group. Size and color relate to the adjusted p-value (pAdj) and log_2_ fold change (L2FC), respectively. (B) Pathway activity of WTAGAAAYY epitope-binding CD8^+^ T cells estimated with PROGENy. Selected significant (Wilcoxon rank sum test) pathways between the conditions are shown. All PROGENy pathways are reported in Supplementary Fig. 5D. (C) Differential transcription factor activity (DoRothEA) estimated with *msviper* in WTAGAAAYY epitope-binding CD8^+^ T cells between severe and mild COVID-19. Positive values indicate increased activity in severe COVID-19. (D) Significantly enriched biological process (BP) gene sets from the Gene Ontology (GO) database and Pathway Interaction Database (PID) gene sets for epitope-binding CD8^+^ T_EM1_ and T_EMRA_ cells. Positive normalized enrichment scores (NES) indicate enrichment in the severe group. TCR = T cell receptor, co-stim = co-stimulatory receptors, EM = effector molecules, IFN = interferon, TF = transcription factor, ISG = interferon-stimulated genes, NKR = NK cell receptors, AP = adaptor proteins, Activ = activation markers, EX = exhaustion markers, CASP = caspase, func = function. *** = p < 0.001, ** = p < 0.01, * = p < 0.05.

For the WTAGAAAYY epitope-binding T_EM1_ and T_EMRA_ cells, genes associated with memory formation were significantly downregulated in severe COVID-19 whereas *PRDM1*, a gene involved in terminal effector T cell differentiation^44^, was overexpressed in T_EM1_ (Fig. 6A). Combined with significant upregulation of caspase 1 and 4, these findings suggested a preferential terminal effector differentiation and a short-lived nature of these epitope-specific CD8^+^ effector populations in severe SARS-CoV-2 infection. However, significantly stronger expression of *CD27* in WTAGAAAYY epitope-binding CD8^+^ T_EMRA_ cells indicated a less differentiated T_EMRA_ phenotype in severe COVID-19 compared to mild infection^45^.

Lower expression of activating and inhibiting NK cell receptors was found for the severe disease group (Fig. 6A). As these receptors have been shown to regulate CD8^+^ T cell effector functions and survival^29,30,34^, this could indicate an impaired regulation of SARS-CoV-2-specific CD8^+^ T cell response and point towards decreased T cell survival in severe disease. We observed significant downregulation of the interleukin 2 receptor beta chain gene (*IL2RB*) in epitope-specific T_EMRA_ cells in severe SARS-CoV-2 infection. Expression of IL2RB is characteristic of a specific memory population which accumulates upon activation of KIR receptors^33^. This finding further supported our hypothesis of impaired CD8^+^ T cell memory formation in severe COVID-19 and was in accordance with our observations from the cell-cell interaction analysis.

Significant upregulation of the activation markers *HLA-DR* and *CD38* together with the RFX transcription factors (Fig. 6C) was observed for the WTAGAAAYY epitope-binding T_EM1_ and T_EMRA_ populations in the severe condition, further underlining hyperactivation of CD8^+^ T cells in severe COVID-19^17^.

Interestingly, analysis of the WTAGAAAYY epitope-specific cells revealed overexpression of exhaustion markers by the T_EMRA_ population in the severe group (Fig. 6A) as an early indication of development towards exhaustion, potentially due to hyperactivation. Lastly, we observed significant negative enrichment of genes associated with “*NFAT TF pathway*” and “*AP1 pathway*” (Fig. 6D) in severe disease. As both NFAT and AP1 are important transcription factors in downstream signaling upon TCR activation^46^, this may indicate impaired signal transduction upon TCR stimulation in severe COVID-19.

In summary, this data suggests that CD8^+^ T cell response to SARS-CoV-2-derived epitopes preferentially leads to short-lived, terminal effector differentiation, and a hyperactivated phenotype in severe COVID-19, while epitope-binding CD8^+^ effector T cells in mild COVID-19 preserve the ability to develop into memory cells after clearance of the viral infection.

## Discussion

While vaccination campaigns are rolled out globally, there is still a limited understanding of the pathogenesis of severe courses of COVID-19. However, this is a critical step in order to develop targeted therapies. A precise understanding of the CD8^+^ T cell response to specific viral epitopes is also important for potential strategies to adjust vaccines in response to the increased occurrence of mutated SARS-CoV-2 variants. Feature barcode technology combined with MHC class I Dextramer reagents carrying specific SARS-CoV-2 epitopes and TCR-seq allowed us to study the epitope-specific CD8^+^ T cell response in COVID-19 at an unprecedented resolution.

We identified different functional subsets of CD8^+^ T cells which seem to acquire an effector phenotype during SARS-CoV-2 infection. However, terminal effector differentiation differed between CD8^+^ T cells in mild and severe disease. While in mild COVID-19, effector cells acquired characteristics indicative of a highly functionally differentiated effector phenotype, CD8^+^ T_EMRA_ cells seemed to be less differentiated in severe disease as indicated by significant overexpression of *CD27*^45^. Furthermore, CD8^+^ effector cells displayed a hyperactivated phenotype in severe COVID-19 and showed significant reduction in characteristics of functional differentiation.

The WTAGAAAYY epitope-specific T_EM1_ and T_EMRA_ populations displayed decreased expression of transcription factors and cytokine receptors associated with T cell memory formation (especially *IL7R*) in severe COVID-19 compared to mild disease (Fig. 6A). Additionally, cell-cell interaction analysis revealed a dysregulation in ligand-receptor interactions associated with long-term T cell memory formation in severe disease. These results might indicate a lower potential for formation of a long-lived SARS-CoV-2-specific CD8^+^ T cell memory in severe COVID-19. Furthermore, the inflammatory environment in severe COVID-19 seemed to favor the development of exhausted CD8^+^ T cells, which might be caused by the hyperactivation of T cells. In line with previous reports^47^, we observed a significant downregulation of IFN-stimulated genes in active severe COVID-19 when compared to active mild. Combined with significantly reduced JAK-STAT signaling and STAT1/2 transcription factor activity in severe COVID-19, this is strongly indicative of an impaired response to IFN stimulation in severe disease.

As previously described, IFN-induced signaling is mediated via STAT1/2, which bind to IFN regulatory factor 9 (*IRF9*) to generate the IFN-stimulated gene factor 3 (ISGF3)^18^. Nuclear transport of ISGF3 is mediated by karyopherin alpha 1 (*KPNA1*) and beta 1 (*KPNB1*)^21,48,49^. It has previously been demonstrated that ORF6 protein from SARS-CoV-1 is able to impair the formation of KPNA1:STAT1:KPNB1 complexes, thereby limiting the IFN response^21^. Furthermore, several mechanisms have been demonstrated by which SARS-CoV-2 impairs the expression and signal transduction of IFN-α and −β^50^. However, in mild COVID-19, JAK-STAT-signaling seems to be sufficient to mount significantly higher expression levels of IFN-stimulated genes compared to severe disease. Interestingly, these findings were reversed when comparing WTAGAAAYY epitope-binding cells between severe and mild COVID-19, indicating that epitope-recognition and TCR stimulation are responsible for these changes.

In addition to TCR activation, IFN-I stimulation is an important signal for T cell activation^51,52^. In non-activated CD8^+^ T cells, type I IFNs induce STAT1/2 signaling to upregulate IFN-stimulated genes. STAT1 has been shown to have strong anti-proliferative effects, thereby preventing expansion of non-specific CD8^+^ T cells^52,53^. Conjugated signaling via IFN-receptors and TCR leads to impaired STAT1 phosphorylation and limits *STAT1* expression^52,53^. Instead, epitope-specific CD8^+^ T cells upregulate *STAT4* expression to transduce IFN-induced signals^51,52^. Through this mechanism, epitope-specific CD8^+^ T cells alter the effect of IFN signaling so that IFN stimulation now promotes proliferation, acquisition of effector functions, and survival via STAT4^51,52^. In our WTAGAAAYY epitope-binding CD8^+^ T cell population we observe upregulation of *STAT1* and downregulation of *STAT4* in severe COVID-19, when compared to mild disease. This might indicate that the necessary upregulation of *STAT4* and downregulation of *STAT1* upon combined TCR and IFN stimulation in epitope-binding CD8^+^ T cells failed in severe SARS-CoV-2 infection. In this case, type I IFNs would still transduce their signals via STAT1, thereby mediating pro-apoptotic and anti-proliferative signals and promoting expression of IFN-stimulated genes^52^, while epitope-binding cells in the mild group effectively downregulate *STAT1* signaling. This would explain the observation of stronger expression of IFN-stimulated genes in WTAGAAAYY epitope-binding CD8^+^ T cells in severe SARS-CoV-2 compared to mild infection. Preferential STAT4 signaling instead of STAT1 in WTAGAAAYY epitope-binding CD8^+^ T cells in mild COVID-19 would also explain the observation of stronger JAK-STAT pathway activity in severe disease as the PROGENy model only includes STAT1-3 and not STAT4.

As NFAT and AP-1 depend on karyopherins for their nuclear translocation^54^, it is possible that SARS-CoV-2-induced inhibition of karyopherin-mediated nuclear transport of STAT1/2 also affects translocation of NFAT in response to TCR activation. This could explain our observation of significantly reduced enrichment of genes associated with “*NFAT TF pathway*” and “*AP1 pathway*” in WTAGAAAYY epitope-binding CD8^+^ T cells in severe disease and on the other hand provide an explanation for impairments in the switch from STAT1 to STAT4 signaling upon combined activation of IFN- and T cell receptor. Impaired nuclear translocation of NFAT upon CD8^+^ T cell activation has been reported as a cause for CD8^+^ T cell exhaustion. In this report, the exhausted T cells were unable to produce cytokines but retained cytotoxic functions^55^. As WTAGAAAYY epitope-binding CD8^+^ T cells display characteristics of exhaustion and strong upregulation of cytotoxic effector molecules at the same time, it is conceivable that the suggested mechanism of impaired nuclear translocation of NFAT contributes to the development of exhaustion.

A prerequisite for the virus to affect karyopherin-mediated nuclear translocation and therefore JAK-STAT signaling is viral entry into CD8^+^ T cells, and the ability to enter immune cells has recently been demonstrated for SARS-CoV-2^56^. We could not detect viral RNA in our scRNA-seq data using Viral-Track^57^. However, limitations are imposed on the ability to detect viral RNA by the tool, including low viral load in immune cells and sparsity of scRNA-seq data^57^. Viral infection of the CD8^+^ T cells could therefore not be ruled out. Weak evidence supporting this hypothesis was the significant enrichment of genes associated with “*viral gene expression*” in the SLEC lineage during severe COVID-19 (Fig. 3D). Assuming viral infection is a contributing factor in observed changes in JAK-STAT signaling and in CD8^+^ T cell differentiation in severe COVID-19, it remains unclear why viral infection leads to severe disease in some individuals, while others only develop mild symptoms.

One possible explanation for the difference in disease progression of COVID-19 could be the significant downregulation of *IFITM3* in severe COVID19. Downregulation of the IFN-stimulated gene was still pertained despite increased JAK-STAT signaling in the WTAGAAAYY epitope-binding T_EM1_ cells in the severe group compared to mild COVID-19. As previously described, IFITM proteins have been implicated in the restriction of SARS-CoV-1 cellular entry^43^ and baseline expression of IFITM3 has been shown to determine the susceptibility to influenza A virus infection^41,42^. It is therefore possible that in individuals who develop only mild COVID-19, a higher baseline expression of IFITM3 restricts cellular entry into CD8^+^ T cells, therefore leading to reduced viral loads and less impairment of JAK-STAT signaling compared to severely infected patients.

The proposal of therapeutic JAK-STAT-inhibition against COVID-19 is gaining popularity^58^. Considering our observations, impaired JAK-STAT signaling seems to be one of the pathomechanisms driving severe COVID-19. Accordingly, therapeutic inhibition of JAK-STAT signaling might aggravate the course of disease. With regard to the observed T cell exhaustion in severe COVID-19, checkpoint inhibition (PD-1 inhibition) could be a potential therapeutic approach as checkpoint inhibition has been suggested to target cells in a pre-exhausted stage and restore their ability to develop into memory T cells^59^.

Epitope-binding analysis of CD8^+^ T cells revealed recognition of eight epitopes from our epitope pool, among them four epitopes derived from the SARS-CoV-2 S protein, three from the N protein, and one from the M protein. Four of these epitopes were uniquely bound by CD8^+^ T cells, with A*0101 WTAGAAAYY showing the largest pool of uniquely responsive CD8^+^ T cells. Interestingly, WTAGAAAYY was found to be the most immunogenic spike protein-derived CD8^+^ T cell epitope according to recent *in silico* predictions^60^.

There are several reasons why we see a reaction against A*0101 WTAGAAAYY in five patients, even though the *in silico* predictions suggested that they are not able to present the epitope on their MHC I molecules. It is conceivable that the reactive CD8^+^ T cells of these five patients actually recognized a different epitope during their infection, which displayed a strong similarity to WTAGAAAYY, accordingly leading to a cross-reactivity with WTAGAAAYY. This assumption could be highly relevant as cross-recognition of WTAGAAAYY by CD8^+^ T cells due to peptide similarities or due to the high immunogenicity of WTAGAAAYY could lead to a broad and unspecific activation of CD8^+^ T cells. Therefore, we identified WTAGAAAYY as a relevant epitope for the generation of CD8^+^ T cell responses to SARS-CoV-2 infection that could be relevant for further development of peptide-based vaccines.

Despite interesting observations, our study had some limitations such as the low number of patients per group. For the analysis of epitope-binding cells, the active and recovered conditions were collapsed to generate an overall group of mildly and severely infected individuals with increased group sizes. Lastly, two of our negative control Dextramer reagents displayed relatively high similarity to peptides derived from other HCoVs (Supplementary Fig. 5E). Since the negative controls need to retain the ability to be presented by MHC class I molecules, this poses limitations to the randomness of their peptide sequences. This necessitated a stringent threshold for specific binding by CD8^+^ T cells which might have led to a loss of signal for certain SARS-CoV-2 derived epitopes.

For the purpose of our study, we aimed to dissect differences between mild and severe COVID-19 by providing an in-depth immunoprofiling of epitope-specific CD8^+^ T cells. We identified WTAGAAAYY as a relevant S protein-derived epitope for the generation of SARS-CoV-2-specific CD8^+^ T cell responses. We demonstrated that CD8^+^ T cells in severely affected patients displayed a dysregulated IFN-response, probably due to impaired JAK-STAT signaling. While CD8^+^ T cells from mildly affected individuals displayed characteristics of highly differentiated effector cells and maintained expression of genes relevant for memory formation, CD8^+^ T cells in severe COVID-19 exhibited a hyperactivated phenotype and differentiated towards exhaustion. Further research has to be conducted to demonstrate viral entry of CD8^+^ T cells as the cause of the observed changes in IFN-response and JAK-STAT signaling, and to investigate whether decreased expression levels of IFITM3 are able to increase the susceptibility of CD8^+^ T cells to viral infection, thereby increasing the risk for a severe course of SARS-CoV-2 infection.

## Acknowledgments

We thank the COVID Aachen Study (COVAS) for contributing patient specimens to the study and the Genomics Facility of the Interdisciplinary Center for Clinical Research (IZKF) Aachen for sequencing experiments. Special thanks goes to Jasmin Hübner and Stephan Klinkenberg for their extraordinary help in sequencing the samples. This work was supported by grants of the German Research Foundation (DFG: KR 4073/11-1; SFBTRR219, Project ID 322900939; CRU344, P1) by a Grant of the European Research Council (ERC-StG 677448), a Grant of the State of North Rhine-Westphalia (Return to NRW) and by BMBF (NUM-COVID19, Organo-Strat 01KX2021) all to RK and DFG(CRU 344, Z) to IC. This work was also supported by the BMBF eMed Consortia Fibromap (to RK, RS and IC). This work was in part supported by the Bioinformatics DATa ENvironment (BioDATEN) project (EF).

## Author Contributions

RK, CK and FS designed the study, FS generated the data and FS, MH, TMA, IC and RK interpreted the data. FS, MH, and RK wrote the manuscript and organized the Fig.s. IK and MB helped with the bulk screening, JK, DF, TMA, GM, NM, MD consented patients and contributed specimens. PZ and TK performed the antibody analysis, and JE, VB and FW performed the HLA typing. IC, RKS, CKur, PB and JSR edited the manuscript and advised on data analysis and interpretation. FS and MH carried out most of the single cell data analysis. FS, MH, JN, and EF carried out cell-cell communication analysis. FS, MH, and FT performed epitope-binding and TCR clonality data analysis. All authors read and approved the final manuscript.

## Methods

### Materials Availability

Epitopes, MHC class I molecules, and DNA-barcode sequences of the used peptide-MHC I Dextramer reagents are provided in Supplementary Table 1. Primer sequences that were used to quantify Dextramer reagent enrichment by bulk sequencing are provided in Supplementary Table 7. There are restrictions to the availability of primer sequences for HLA typing as these primers were developed for the use in clinical practice at the University Hospital Düsseldorf, Düsseldorf, Germany. These primers are still used by the Institute of Transplantation Diagnostics and Cell Therapeutics (ITZ), University Hospital of Düsseldorf, Düsseldorf, Germany. Accordingly, the sequences are subject to intellectual property protection.

### Patient recruitment and clinical data

25 laboratory-confirmed COVID-19 patients were recruited from the University Hospital of the RWTH Aachen University and from the Sankt Antonius Hospital Eschweiler from May to September 2020. 7 healthy volunteers were included in the study. All patients provided informed consent and the study was performed in accordance with the Declaration of Helsinki. For patients who were not able to give consent themselves, their legal representative agreed to their participation in the study. The study protocol was reviewed and approved by the Ethical Board of the RWTH Aachen University Hospital (vote: EK 078/20). Three COVID-19 patients and one healthy volunteer were excluded from the study due to missing clinical data and bad sample quality, respectively. Thus, 22 COVID-19 patients and 6 volunteers were included in the study. Patients were pseudonymized and clinical as well as epidemiological data were obtained from the electronic hospital information system “CGM Medico”. Clinical data is provided in Supplementary Table 2.

### Group allocation

Based on the clinical course and time point of blood withdrawal, patients were divided into two major groups; severe or mild SARS-CoV-2 infection. Patients with asymptomatic infection and symptomatic patients who did not require mechanical ventilation were allocated to the group of mild infection. Symptomatic patients who required mechanical ventilation were allocated to the group of severe SARS-CoV-2 infection.

Based on the PCR result that was closest to blood withdrawal, patients within the major groups of mild or severe infection were further subdivided into the subgroups “active infection” or “recovered”. If a PCR result of a previously positive localization was negative before or at the time of blood collection and the result subsequently remained negative as well, the patients were assigned to the group of recovered SARS-CoV-2 infections. If a positive PCR result was obtained on the day of blood collection or later, or if the last positive PCR result was obtained at a maximum of 3 days before blood collection, the patients were assigned to the group of active SARS-CoV-2 infections. Group allocation for each patient is shown in Supplementary Table 2.

### Sample collection and PBMC isolation

10-30 ml of blood per patient were collected in either 9 ml S-Monovettes, K3 EDTA 92×16 mm, or in 5.5 ml S-Monovettes, 75×15 mm, provided by Sarstedt (Nümbrecht, Germany). The samples were stored at 4°C for a maximum of 7 hours until further processing. Peripheral blood mononuclear cells (PBMCs) were isolated by Ficoll gradient centrifugation. The isolated PBMCs were resuspended in 10% DMSO in FCS and immediately frozen gradually. The frozen PBMC samples were stored at −152°C. In addition, serum samples were frozen for each patient. After thawing, the PBMCs were immediately diluted with 10 ml 5% FCS in PBS and centrifuged at 500 rcf for 5 minutes. The supernatant was removed and the PBMCs were resuspended in 5 ml 5% FCS in PBS and filtered through a 20 μm pluriStrainer**®** provided by pluriSelect (Leipzig, Germany) to obtain a single cell solution.

### SARS-CoV-2 antibody testing

To exclude healthy controls with a previous SARS-CoV-2 infection, we tested the subjects for SARS-CoV-2-specific IgG antibodies by performing the Euroimmun anti-SARS-CoV-2 ELISA (IgG) (EUROIMMUN, Lübeck, Germany) (Supplementary Fig. 1D). An IgG ratio of > 2.5 was considered a positive test, a ratio between 0.8 and 2.5 was considered an intermediate result and a ratio < 0.8 was considered a negative test^61^.

### High-throughput amplicon-based HLA typing

DNA for HLA typing was isolated from PBMCs using the Quick-DNA/RNA™ Microprep Plus Kit (Zymo Research, Irvine, CA, USA). High throughput-HLA typing was performed by the Institute of Transplantation Diagnostics and Cell Therapeutics (ITZ), University Hospital of Düsseldorf, Düsseldorf, Germany using an amplicon based Next Generation Sequencing assay that targets the HLA genes HLA-A, -B, -C, -DRB1, -DQB1 and -DPB1.

Primers for HLA typing were designed to target exons 2 to 4 for the HLA class 1 genes as well as for HLA-DPB1 and exons 2 and 3 for HLA-DRB1 and HLA-DQB1. Amplificates comprised the entire exon each and additional flanking intron sequences. Primers were screened for SNPs using the SNPCheck software (https://genetools.org/SNPCheck/snpcheck.htm) to prevent impaired primer binding and subsequent allele drop-out and erroneous genotyping. Primers were purchased from Biolegio (Nijmegen, The Netherlands). Each primer pair was checked for specificity using Sanger sequencing. For intellectual property reasons, primer sequences that are used in the Next Generation Sequencing assay for HLA typing are not described here.

### Next Generation Sequencing and workflow for HLA typing

The entire set of fragments was amplified in six multiplex PCR reactions. After a clean-up step using paramagnetic beads, sample-specific barcodes and Illumina compatible adapter sequences were added in a second-round PCR. Samples obtained from the UCLA International Cell Exchange program were included as quality controls for both PCR steps. The samples were pooled, underwent a second purification step, and were quantified using the QuantiFluor dsDNA system (Promega, Walldorf, Germany). Paired-end sequencing was performed on a MiSeq platform (Illumina, San Diego, CA, USA) for 7 pM of the library with 2x 280 cycles using a standard v3 cartridge. As an internal quality run control, we used a spike-in of 15% of PhiX DNA. After de-multiplexing using the MiSeq Reporter software (Illumina Inc.), the analysis of the read sequences was performed by a Visual Basic-based in-house software (NGSSequence Analyser, Institute of Transplantation Diagnostics and Cell Therapeutics (ITZ), University Hospital of Düsseldorf, Düsseldorf, Germany) approach considering quality control values and high coverage to automate data analysis.

Sequencing runs had to pass the following criteria: cluster density between 800 and 1,300 k/mm2, more than 70% of clusters passed the filter, Q30 score must be more than 70%.

MHC class I molecules show overlapping peptide specificity as they bind the same main anchor motifs in peptides^62^. According to this shared peptide recognition, MHC-class I molecules can be grouped into so-called HLA-supertypes^62^. Therefore, patients will likely not only be able to present an epitope if they match the exact HLA-allele of a certain Dextramer reagent, but also if their HLA-allele matches the HLA-supertype of the Dextramer reagent. To compare the HLA-supertypes between patients and Dextramer reagents, we determined the HLA-supertype for the HLA-A and HLA-B alleles. HLA-alleles and the associated HLA-supertypes are shown in Supplementary Table 3.

### SARS-CoV-2 epitope selection and synthesis of MHC I Dextramer reagents

An explorative panel of 30 SARS-CoV-2-derived epitopes with the strongest predicted affinity to MHC class I molecules was defined by Immudex (Copenhagen, Denmark). The selection was based on internal predictions using NetMHCpan version 4.0 and on previous publications, followed by confirmed binding of the selected epitopes to the assigned MHC I molecules. Synthesis of the peptide-MHC class I Dextramer reagents for both, bulk and single-cell experiment, was performed by Immudex. Dextramer reagents consisted of a dextran backbone to which several MHC-I molecules were attached, each of them carrying one of the selected SARS-CoV-2 epitopes. Each Dextramer reagent carried a specific DNA barcode, and a PE-label. As the MHC molecules of one MHC I Dextramer reagent carried the same SARS-CoV-2 epitope, subsequent identification of the epitopes was possible using the Dextramer reagent-specific DNA barcode. Additionally, 8 control dCODE Dextramer reagents were included into the panel. Dextramer reagents carrying nonsense peptides were used as negative controls and Dextramer reagents carrying CMV-derived epitopes were used as positive controls. An overview over the epitope panel is provided in Supplementary Table 1.

### Quantification of epitope-specific T cell responses in a bulk-approach

PBMC samples were thawed as described above. After the PBMCs were passed through a strainer, a second centrifugation was performed. Samples were enriched for T cells by magnetic depletion of the remaining blood cell fractions from the PBMC layer using the Pan T cell Isolation kit, human (Miltenyi Biotec). The dCODE Dextramer reagents (HiT) were pooled, using 2 μl of each Dextramer reagent per patient. In accordance with the protocol provided by Immudex, 0.2 μl 100 μM d-Biotin were added per Dextramer reagent specificity. T cells were resuspended in 100 μl 5% FCS in PBS including Herring sperm DNA at a concentration of 0,1 g/l and subsequently incubated with the pool of Dextramer reagents. Excess Dextramer reagents were removed by several washing steps. Using magnetic activated cell sorting and Anti-PE MicroBeads (Miltenyi Biotec), we separated the T cells into a Dextramer reagent-positive (PE^+^) and Dextramer reagent-negative (PE^−^) fraction.

The PE^+^ and PE^−^ fractions were processed identically in a two-step PCR procedure using KAPA HiFi HotStart ReadyMix (Roche). The protocol for the two-step PCR approach was adopted and modified from a previously described method^63^. During the first PCR, the DNA barcodes of the MHC I Dextramer reagents were amplified using Dextramer reagent-specific primers, which contained an overhang that functioned as a general handle sequence for the primers in the second PCR (Supplementary Table 7). The nucleotide sequence for the general handle sequences were adopted from the protocol described by Franzen et al^63^. To generate a single-indexed Illumina-compatible library, a second PCR was performed. As a reverse primer we used custom “barcoded primers” including the Illumina i7 adapter sequences, an 8bp sample index and sequences that were complementary to the general handle sequence from the first PCR. As a forward primer we used a “common primer” including only the Illumina i5 adapter sequences and the sequences that were complementary to the general handle sequence from the first PCR (Supplementary Table 7). Nucleotide sequences of SureSelect^XT^ Indexes from the “SureSelect^XT^ Target Enrichment System for Illumina Multiplexed Sequencing” were included as 8bp sample indexes in the “barcoded primer”. All custom PCR primers were purchased from Eurofins Genomics (Luxembourg). Primer sequences are provided in Supplementary Table 7. After every PCR, PCR products were purified using the QIAquick^®^ PCR Purification Kit (Qiagen). Cycling conditions for the first PCR were as follows: Initial denaturation 98°C for 45 sec, denaturation 98°C for 15 sec, annealing 60°C for 30 sec, extension 72°C for 30 sec and final extension 72°C for 1 minute; denaturation, annealing and extension were repeated for a total of 16 cycles. Cycling conditions for the second PCR were as follows: Initial denaturation 98°C for 45 sec, denaturation 98°C for 15 sec, annealing and extension 72°C for 30 sec and final extension 72°C for 1 minute; denaturation, annealing and extension were repeated for a total of 16 cycles.

The Dextramer reagent-positive and -negative samples were pooled equimolar after fluorometric quantification. The library pool was quantified by qPCR and sequenced on a MiSeq platform (Illumina, San Diego, CA, USA) with 2x 150 cycles. To compensate for the low complexity of the amplicon library, up to 50% phiX DNA was added.

### Bulk sequencing data analysis

Bulk-seq data analysis was carried out using the *ShortRead* package^64^ in *R* version 3.6.3. Sequencing data for the Dextramer reagent-positive and -negative sample were processed separately. After extraction of unique reads to avoid amplification bias, the number of barcodes for each MHC I Dextramer reagent was calculated in the samples. According to the workflow provided by Immudex, the apparent enrichment (AE) was calculated by dividing the counts for every Dextramer reagent in the Dextramer reagent-positive sample by the counts in the negative samples. For the negative controls, a median AE was calculated. Finally, the specific enrichment was calculated by dividing the AE for every Dextramer reagent by the median AE of the negative controls. Every Dextramer reagent that displayed a specific enrichment value of at least 5 in at least one condition was carried into the single-cell analysis. Based on this approach, we finally selected 15 Dextramer reagents and the 8 control Dextramer reagents for the single-cell experiment. To select three patients for each condition for the single-cell experiment, we divided Dextramer reagent-enrichment in the bulk experiment into four groups, based on the comparison between the HLA alleles of the patients and the Dextramer reagents and the specific enrichment value. HLA-match and a log_2_-fold enrichment < 1 was considered “no antigen specificity”. No HLA-match and log_2_-fold enrichment < 1 was considered “not HLA compatible”. HLA-match and log_2_-fold enrichment ≥ 1 was considered a “specific enrichment”. No HLA-match and log_2_-fold enrichment ≥ 1 was considered “unspecific binding”. For the single-cell experiment, we finally chose the three individuals from every condition, who displayed the most “specific enrichment”.

### Single-cell immune profiling of epitope-specific T cells

PBMC samples were thawed as described above. After the PBMCs were passed through a strainer, a second centrifugation was performed. The supernatant was removed and cells were diluted in 100 μl 5% FCS in PBS. 5 μl Human TruStain FcX (BioLegend) was added and incubated for 10 minutes at 4°C. The panel of 23 10x-compatible MHC class I Dextramer reagents (Immudex), including 8 control Dextramer reagents was pooled as described above. Furthermore a panel of 15 TotalSeq-C antibodies (BioLegend) was pooled, using 0,5 μg of each antibody. The Dextramer reagent pool was incubated with the samples for 10 minutes at 4°C, followed by a 30 minute incubation with the TotalSeq-C antibody pool and 5 μl of a PE/Cyanine7 anti-human CD8 antibody (BioLegend) at 4°C. Cells were washed two times using 3 ml 5% FCS in PBS and rediluted in 2 ml 5% FCS in PBS. For the detection of dead cells, DAPI was added at a final concentration of 0,5 μg/ml. Cells were sorted into 1% BSA in PBS on a Sony Cell Sorter by gating on two populations; a PE^+^ population (MHC I Dextramer reagent-positive) and a CD8^+^ PE^−^ fraction (Supplemental Fig. 2A). If the number of PE^+^ cells did not exceed 10.000 cells, a maximum of 3.000 CD8^+^ PE^−^ cells were added prior to single-cell partitioning and barcoding via Chromium Controller (10x Genomics). For each sample, three libraries were prepared; a 5’ gene expression library (GEX), a T cell receptor enriched library (VDJ), and a surface protein library containing the TotalSeq-C and MHC I Dextramer reagent barcodes (ADT). After fluorometric quantification, the libraries were pooled in a 5:1:1 ratio for the GEX library, the VDJ enriched library and the ADT library, respectively. The pooled libraries were quantified by qPCR and sequenced on a NextSeq 500 platform (Illumina, San Diego, CA, USA) with 2x 150 cycles.

### Single-cell RNA seq data processing

Raw scRNA-seq FASTQ files were aligned to the human GRCh38 genome with *Cell Ranger* version 4.0.0 with default settings (10x Genomics). For every patient, the paired GEX and ADT libraries were processed together with the *count* function and the VDJ enriched library was processed separately with the *vdj* method. Downstream analysis was conducted with *Seurat* version 4.0^65^ in *R* version 4.0.3. Cells with < 200 or > 3,000 detected genes and more than 10% mitochondrial read content were filtered out (Supplementary Fig. 2G is referred to for scRNA quality control metrics per sample). The GEX and ADT assays were log and centered log ratio (CLR) normalized, respectively, and were subsequently scaled with default settings.

### Clustering and cell annotation

In order to cluster and characterize cell subtypes, the samples were integrated based on the GEX libraries for a first round of clustering. For every dataset, the top 2.000 most variable genes were determined and dimensional reduction was performed on the variable features with a principal component analysis (PCA). The samples were integrated using the *harmony* algorithm^66^ with default settings and the data was embedded in a Uniform Manifold Approximation and Projection (UMAP) using 30 principal components. A shared nearest neighbor graph was built with 30 principal components using *FindNeighbors* and unsupervised clustering was performed using a Louvain-based algorithm with *FindClusters* and a resolution of 1. In order to determine cluster-specific markers, a Wilcoxon rank sum test was performed with *FindMarkers* using min.pct = 0.25. Only genes with a false discovery rate (FDR) < 5% were considered. High-level cell annotation of the clusters was performed on the integrated data followed by filtering of non-T cells and clusters consisting mainly of low-quality cells. A second round of clustering was performed as described above using a resolution of 0.5. Low-level annotation of the resulting clusters was based on a combination of GEX and ADT marker expression as listed in Supplementary Table 4. One healthy sample was removed from the study as the volunteer informed us of an unknown infection in early January 2020 and exhibited high levels of differentiated effector T cells (Supplementary Fig. 2E-F is referred to for patient and condition projection onto the integrated UMAP). Cell-cycle analysis was performed on the clusters using *CellCycleScoring* (Supplementary Fig. 2B) and mitochondrial gene content was computed per disease condition (Supplementary Fig. 2H).

### Differential gene expression and gene set enrichment analysis

For functional characterisation of the differences between the disease conditions, differential gene expression analysis was performed with *FindMarkers* using min.pct = 0.25 and FDR < 5%. When contrasting conditions, only cell types with counts > 20 in both groups were considered for the analysis. A pre-ranked gene set enrichment analysis was performed with the *fgsea* package^67^. The gene sets C2, C5 (subcategory BP), and C7 were used for the analysis and were downloaded with the *msigdbr* package. The top enriched gene sets with an FDR < 5% were visualized as bar plots. A full list of significantly differentially expressed genes for each test is listed in Supplementary Table 5.

### Signaling pathway and transcription factor activity

Signaling pathway activities were estimated with PROGENy^68,69^ using the top 500 footprint genes per pathway. To test for significant differences between the active mild and severe condition, Wilcoxon rank sum tests were performed on relevant pathways and cell types with FDR < 5%. Transcription factor activities were computed with *msviper*^70^ using regulons with confidence levels A, B or C from DoRothEA^71^. The changes in transcription factor activity were estimated per cell type using the condition contrasts obtained from differential gene expression analysis with *FindMarkers*. Transcription factor activities with FDR < 5% or an absolute normalized enrichment score > 2.5 between the active mild and severe condition were visualized with *pheatmap*. Supplementary Fig. 3A is referred to for all PROGENy pathway activities per condition.

### Cell-cell communication

To estimate cell-cell interactions between the CD8^+^ T cell populations, CellPhoneDB version 2.1.5^72^ was used per disease condition with the *statistical_analysis* method. Cell types with counts > 20 per condition were included in the analysis and the log-normalized and scaled counts were used as input. Ligand-receptor interactions with p-value < 0.05 were considered for the downstream analyses. Summarized ligand-receptor interactions between cell types were visualized with *pheatmap* (Supplementary Fig. 3B). For the active mild and severe conditions, interactions with the T cell activating (NKG2D, NKG2C:CD94, CD94:NKG2E, and CLEC2B)^31,73,74^ and inhibiting receptors (NKG2A:CD94, KIR3DL2, KIR2DL3, KIR2DL1, and KLRB1)^75^ were visualized with the *circlize* package^76^.

CrossTalkeR^77^ was used to compute changes in ligand-receptor interactions between the conditions. Briefly, CrossTalkeR constructs representations of the ligand-receptor networks for each condition, where the edges of the network are weighted by the number of interactions and the sum of weights of the interaction-pairs obtained by CellPhoneDB. Differential cell-cell interaction networks were constructed by subtracting the condition state network from the control states. Differential SELL - SELPLG interactions between the recovered mild and severe groups were visualized as sankey plots (Fig. 2G). Sankey plots of differential MICB - NKG2D, HLA-E - CD94:NKG2A, and SELL - SELPLG interactions were also made between the active mild and severe conditions (Supplementary Fig. 3D).

### Trajectory inference and pseudotemporal differential gene expression

In order to estimate activation trajectories of the integrated scRNA-seq data, trajectory inference was performed with *Slingshot*^78^. The MAIT, ɣδ, NKT, and CD8^+^ CD73^+^ T_reg_ populations were excluded from the pseudotime analysis in order to only include T cell subpopulations likely to originate from the CD8^+^ T_N_ cells. *Slingshot* was run on the UMAP embedding of the remaining clusters and the CD8^+^ T_N_ population was designated as the root. Two trajectories were determined by the pseudotime analysis; the short-lived effector cell (SLEC) lineage and memory-precursor effector cell (MPEC) lineage. In order to test for significant differences in the distribution of the active mild and severe conditions across pseudotime, a Kolmogorov-Smirnof test was performed for each lineage.

For temporal differential gene expression analysis between the two trajectories, *tradeSeq* was used^79^. A negative binomial generalized additive model (NB-GAM) was built on the 10,000 most variables genes and pseudotimes for the active mild and severe conditions using the *fitGAM* function. In order to study differences in temporal gene expression between the conditions, a condition-specific smoother was computed per lineage. 6 knots were used for the NB-GAM (Supplementary Fig. 4D is referred to for a visualization of the knots projected onto the integrated UMAP). Differential gene expression between the progenitor and differentiated cell populations was performed with *startVsEndTest* using l2fc = log_2_(2). The significant genes from the test were modeled with *predictSmooth* using nPoints = 50 and visualized with *pheatmap*. The expression of significant genes across pseudotime was visualized with *plotSmoothers*. To characterize potential early drivers of differentiation towards the two trajectories, *earlyDETest* was used at the bifurcation point (between knots 2 and 3) with l2fc = log_2_(1.5). Differential gene expression between the end stages of the lineages was performed with *diffEndTest* using l2fc = log_2_(2). Temporal differential expression between the active mild and severe conditions for each lineage was computed with *conditionTest* using l2fc = log_2_(2), global = TRUE, and pairwise = TRUE. For all tests performed with *tradeSeq*, only genes with FDR < 5% were considered. For each test performed with *tradeSeq*, all genes were ranked based on the estimated Wald statistic and a gene set enrichment analysis was performed as previously described. Supplementary Table 5 is referred to for a full list of significantly differentially expressed genes for each test.

### T cell receptor clonality analysis

In order to study T cell receptor (TCR) clonality in the scRNA data, TCR clonotypes were assigned based on the VDJ library using the *cellranger vdj* function. For the analysis, only MHC class I restricted T cell subtypes were considered. The clonotypes were grouped based on the level of expansion and designated as; single (*n* = 1), small (1 < *n* ≤ 5), medium (5 < *n* ≤ 20), large (20 < *n* ≤ 100), or hyperexpanded (*n* > 100). The relative abundance of the clonotype size groups was computed for the conditions and cell types and visualized as bar charts. Supplementary Fig. 5A is referred to for the TCR clonotype size distribution for each cell type per condition.

The similarity between the MHC class I restricted CD8^+^ T cell subtypes was calculated as the Morisita-Horn overlap of the TCR clonotypes. In order to estimate changes in TCR diversity between the disease conditions, the relative richness and evenness were calculated over pseudotime for the SLEC and MPEC trajectories. To increase the number of samples for the severe and mild conditions for the downstream analysis, the active and recovered groups were collapsed. The TCR richness of the mild and severe conditions was defined as the number of unique clonotypes divided by the total number of cells with an assigned clonotype^80^. TCR evenness was calculated as the inverse Simpson index divided by the number of unique clonotypes^81^.

### SARS-CoV-2 epitope binding of CD8^+^ T cells

To characterise SARS-CoV-2 epitope-specific CD8^+^ T cells, the ADT library of the MHC I Dextramer reagents carrying SARS-CoV-2-derived peptides was used. We followed the protocol from 10x Genomics and Immudex for the identification of epitope-binding single T cells (https://www.10xgenomics.com/resources/application-notes/a-new-way-of-exploring-immunity-linking-highly-multiplexed-antigen-recognition-to-immune-repertoire-and-phenotype/). First, the log_2_ fold-change between the expression of the Dextramer reagents and the median of the negative controls (*n* = 4) was calculated. For every Dextramer reagent, cells were classified as epitope-binding if at least 30% of the cells belonging to the same clonotype had a log_2_ fold change > 2 and had a clonotype expansion size > 5. Epitope binding was found for eight Dextramer reagents, of which four had uniquely binding cells (Supplementary Table 6). Unique epitope-binding counts for the healthy, mild, and severe conditions were visualised as bar charts. Supplementary Fig. 5B is referred to for unique binding counts per patient. Unique binding of the A0101 WTAGAAAYY epitope was shown for the most hyperexpanded TCR clonotypes on the integrated UMAP (Supplementary Fig. 5C).

Clonal expansion of the uniquely WTAGAAAYY-binding cells across pseudotime were visualized as scatter plots for the SLEC and MPEC lineages using *stat_smooth* (method = loess) from the *ggplot2* package. For the uniquely epitope-specific CD8^+^ T cells, the frequency of TRA and TRB CDR3 usage was computed for the mild and severe conditions and the top 15 CDR3 sequences were visualized as bar charts.

In order to dissect, whether the individuals binding to A*0101 WTAGAAAYY in our experiment would be able to present the epitope on endogenous MHC I molecules, we predicted the binding of WTAGAAAYY to HLA alleles in our patient population using NetMHCpan version 4.1 with default settings^82^.

### Functional characterization of antigen-specific CD8^+^ T cells

For a robust characterization of the differences in SARS-CoV-2 specific CD8^+^ T cells between the mild and severe conditions, we focused on subclusters with > 200 uniquely WTAGAAAYY epitope-binding cells with counts originating from at least two patients (Supplementary Fig. 5B). The T_EMRA_ and T_EM1_ populations matched these criteria and were used for the downstream analysis. Differential expression between the mild and severe conditions for each cell type was performed with *FindMarkers*, as previously described. Gene set enrichment analysis together with estimation of pathway signaling and transcription factor activity was computed as described above.

### Protein conservation of viral-epitope panel

To understand possible cross-reactivity of T cells to certain SARS-CoV-2-derived epitopes, protein similarities between the Dextramer reagents and seven homologous human coronaviruses were computed. Multiple sequence alignment was performed with the *msaClustalOmega* function from the *msa* package^83^. Distance matrices of the aligned sequences were computed with *dist.alignment* (matrix = identity) from the *seqinr* package^84^ and the pairwise protein similarities were defined as 1 - distance. The protein conservation of the Dextramer reagents were visualized as tile plots. Supplementary Fig. 5E is referred to for protein sequence similarities between the negative Dextramer controls used in this study (*n* = 4) and Spike proteins of known human coronaviruses.

### SARS-CoV-2 viral detection

For the quantification of SARS-CoV-2-derived viral reads in the scRNA data, *Viral-Track* was used^57^. The viral genome of the German SARS-CoV-2 isolate (accession MT270101) was added to the human GRCh38 genome and *STAR* (runmode genomeGenerate)^85^ was run to create a joint reference genome. The *Viral-Track* pipeline was subsequently run on the FASTQ files of each sample with default settings.

