## Supplemental Figures and Tables for "Dissecting CD8+ T cell pathology of severe SARS-CoV-2 infection by single-cell epitope mapping"

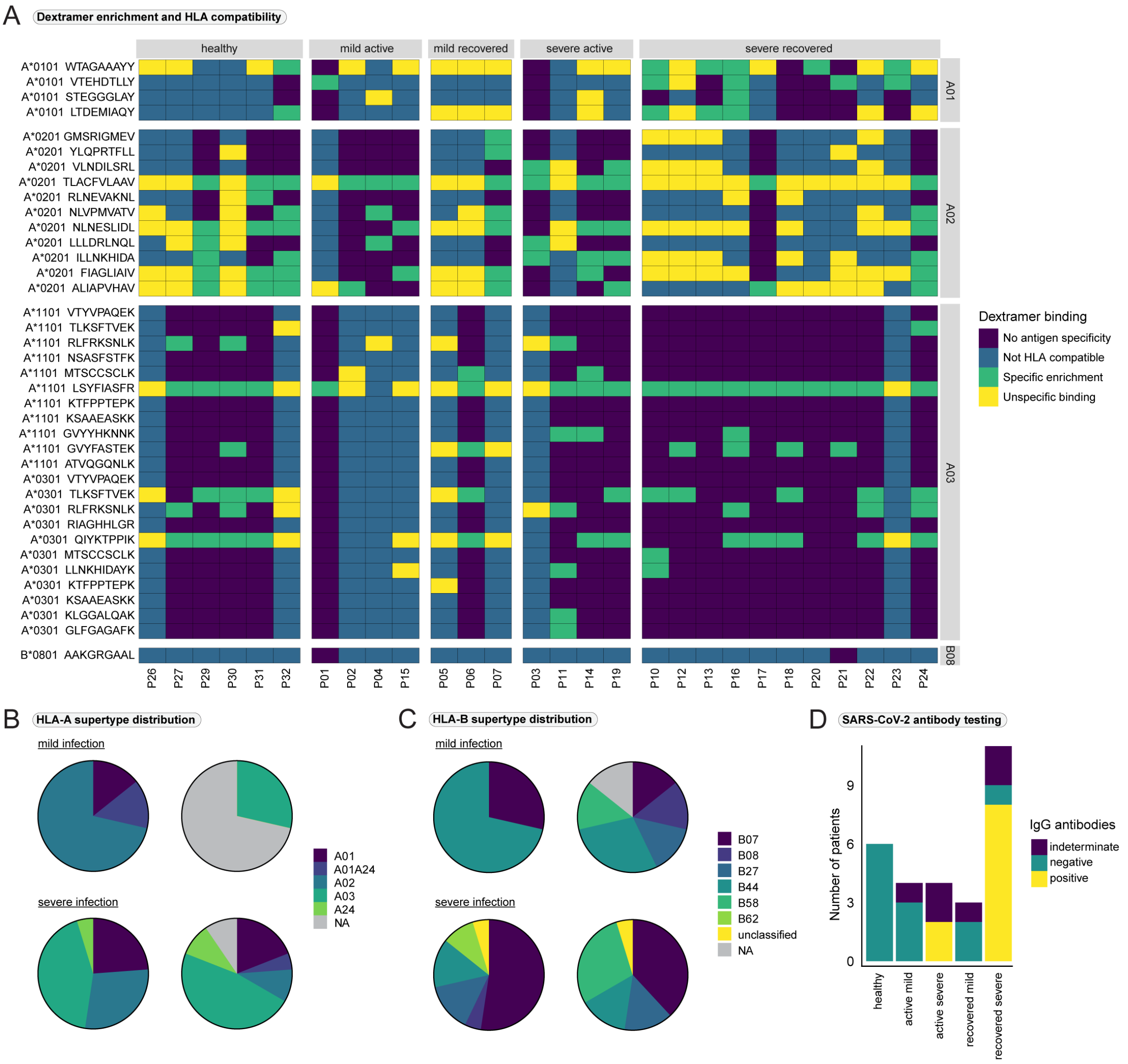


**Supplemental Fig. 1. Overview of patient and Dextramer reagent HLA-alleles, Related to Fig. 1 and Supplemental Table 1 and 3. (A)** Classification of epitope-binding characteristics in the bulk screening experiment according to the specific enrichment value and the match between HLA supertypes of the patients and Dextramer reagents. Patients are displayed on the x-axis, SARS-CoV-2 epitopes and the corresponding HLA alleles are displayed on the y-axis. No antigen specificity = HLA-match and $log_2$-fold enrichment < 1. Not HLA compatible = no HLA match and $log_2$-fold enrichment < 1. Specific enrichment = HLA match and $log_2$-fold enrichment ≥ 1. Unspecific binding = no HLA match and $log_2$-fold enrichment ≥ 1. **(B)** Overview of the distribution of HLA-A supertypes per overall condition (mild and severe disease). For each condition, HLA-A allele 1 (left) and HLA-A allele 2 (right) is depicted. **(C)** As in (B) but for the HLA-B supertype. **(D)** Results of SARS-CoV-2 IgG antibody testing.

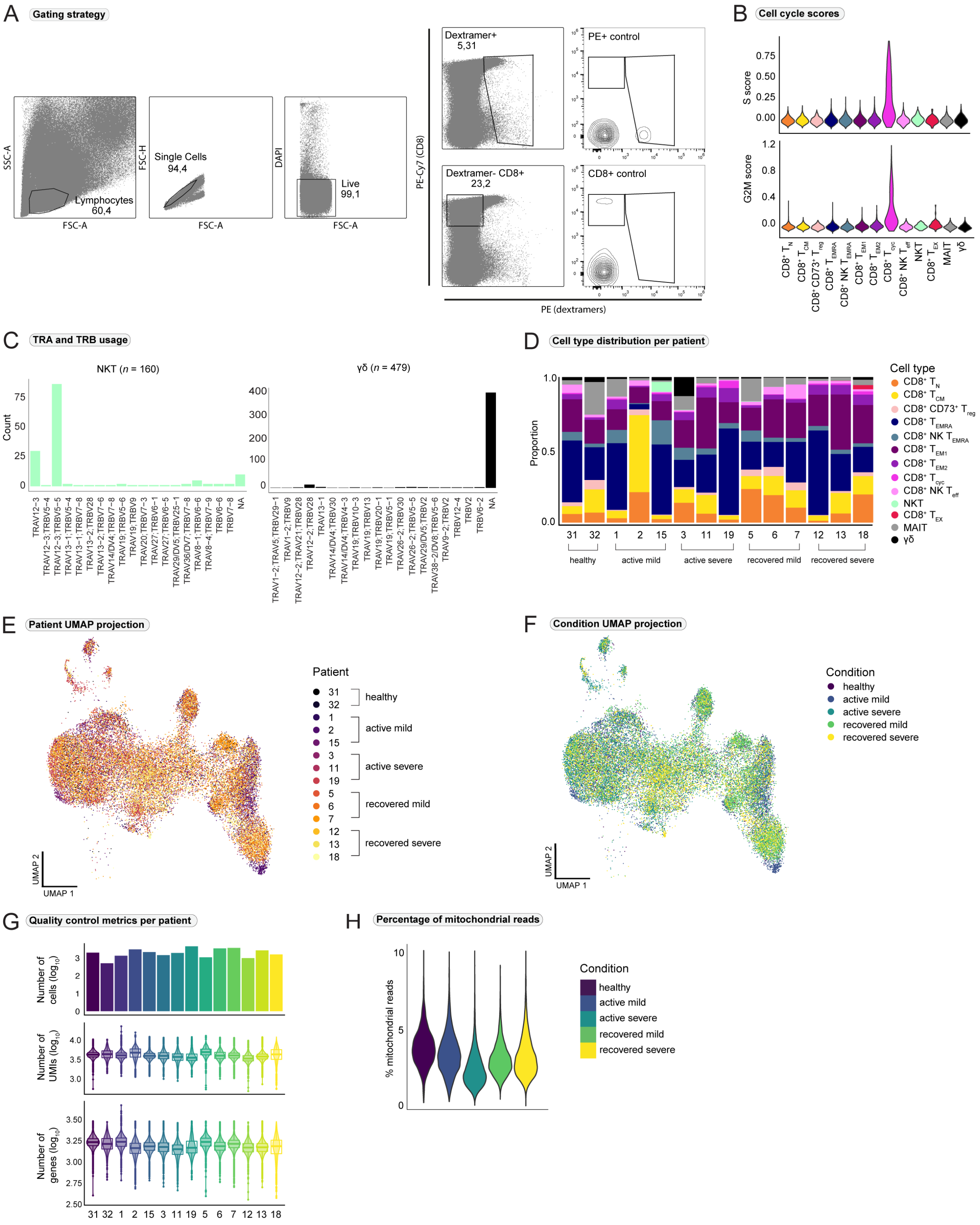


**Supplemental Fig. 2. Additional information on the generation of single-cell data, Related to Fig. 1. (A)** FACS gating strategy. Two different populations were sorted for, a $PE^+$ (Dextramer reagent-positive) and a $CD8^+$ $PE^-$ population. **(B)** Cell cycle scores per $CD8^+$ T cell subtype. **(C)** T cell receptor alpha (TRA) and T cell receptor beta chain (TRB) usage in atypical NKT cells and γδ T cells. **(D)** Average distribution of $CD8^+$ T cell subsets per patient. **(E)** Per patient-origin and **(F)** per condition-origin projected onto the integrated UMAP. **(G)** Overall single-cell RNA-seq data quality control metrics per patient. **(H)** Percentage of mitochondrial read content per condition.

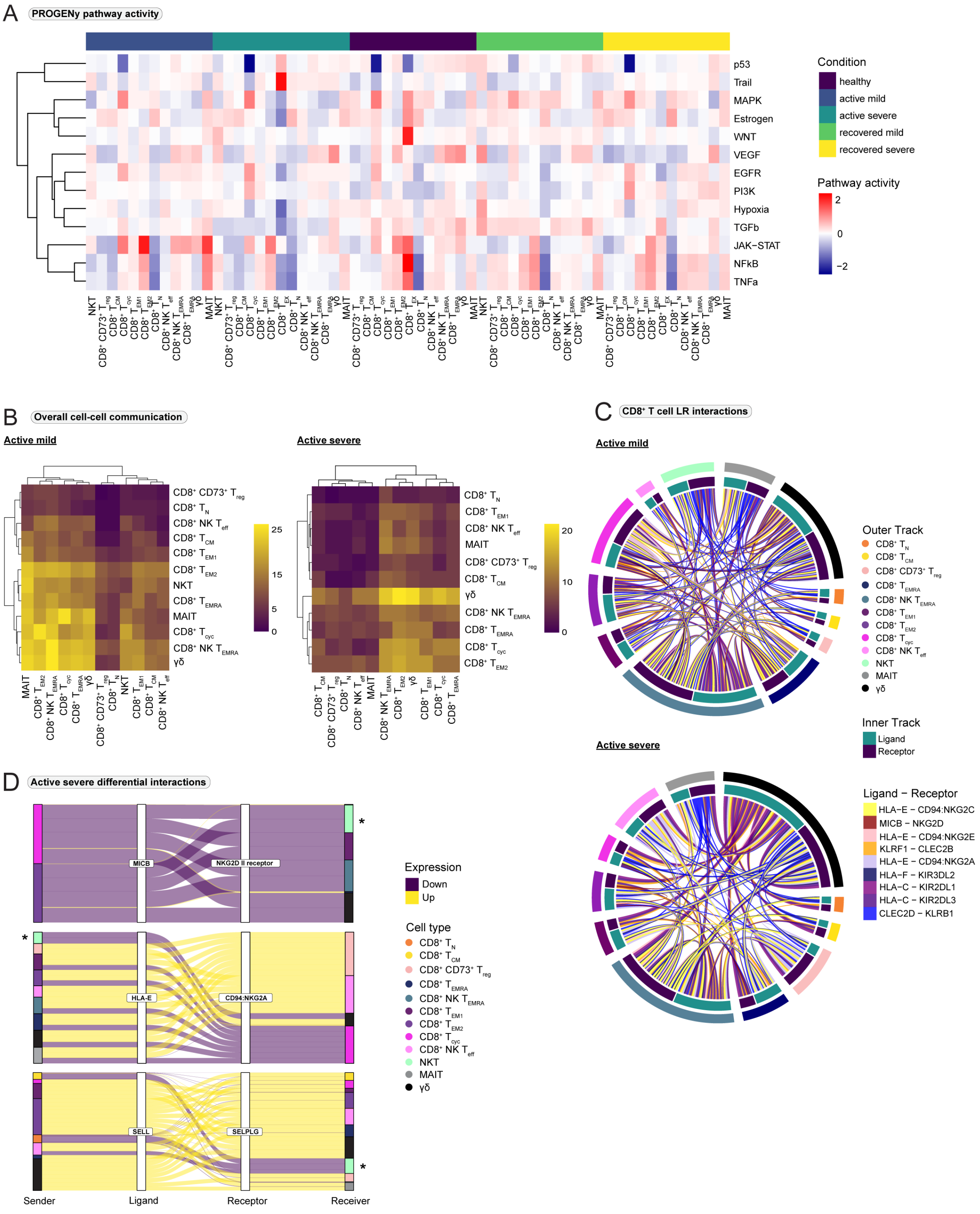


**Supplemental Fig. 3. Pathway analysis and cell-cell interactions, Related to Fig. 2. (A)** Pathway activity for all computed PROGENy pathways. **(B)** Overall frequency of cell-cell interactions in active mild (left) and severe COVID-19 (right). Senders (ligands) are displayed on the x-axis and receivers (receptors) are displayed on the y-axis. **(C)** Selected T cell activating (warm colors) and inhibiting (cold colors) ligand-receptor (LR) interactions between $CD8^+$ T cell subtypes. **(D)** Sankey plot of differential interactions of MICB - NKG2D, HLA-E - CD94:NKG2A, and SELL - SELPLG between active mild and severe COVID-19. Positive values indicate increased interactions in the active severe group. *NKT cells were only included for the active mild condition.

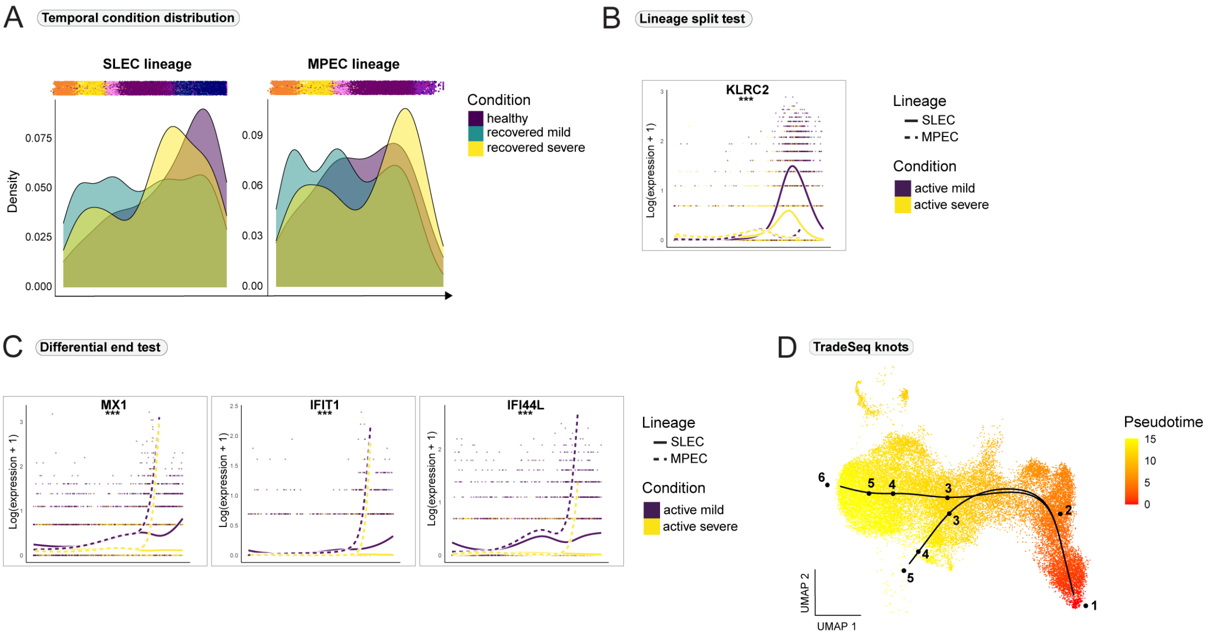


**Supplemental Fig. 4. Relevant genes in CD8$^+$ T cell differentiation during SARS-CoV-2 infection identified by tradeSeq, Related to Fig. 3 and Supplemental Table 5. (A)** Temporal distribution of cell density per condition and trajectory. Fig. 3B is referred to for the active mild and severe conditions. **(B)** Expression of the gene *KLRC2* that was predicted to be significantly differentially expressed at the bifurcation of the two lineages. Differential expression was estimated with *tradeSeq*. The y-axis is on the natural logarithmic scale. **(C)** Significantly differentially expressed genes between the two lineages in the end stages of CD8$^+$ T cell differentiation. The y-axis is on the natural logarithmic scale. **(D)** *tradeSeq* knots that subdivide pseudotime into partitions, projected onto the integrated UMAP. SLEC = short-lived effector cells, MPEC = memory precursor effector cells. *** = p-value < 0.001.

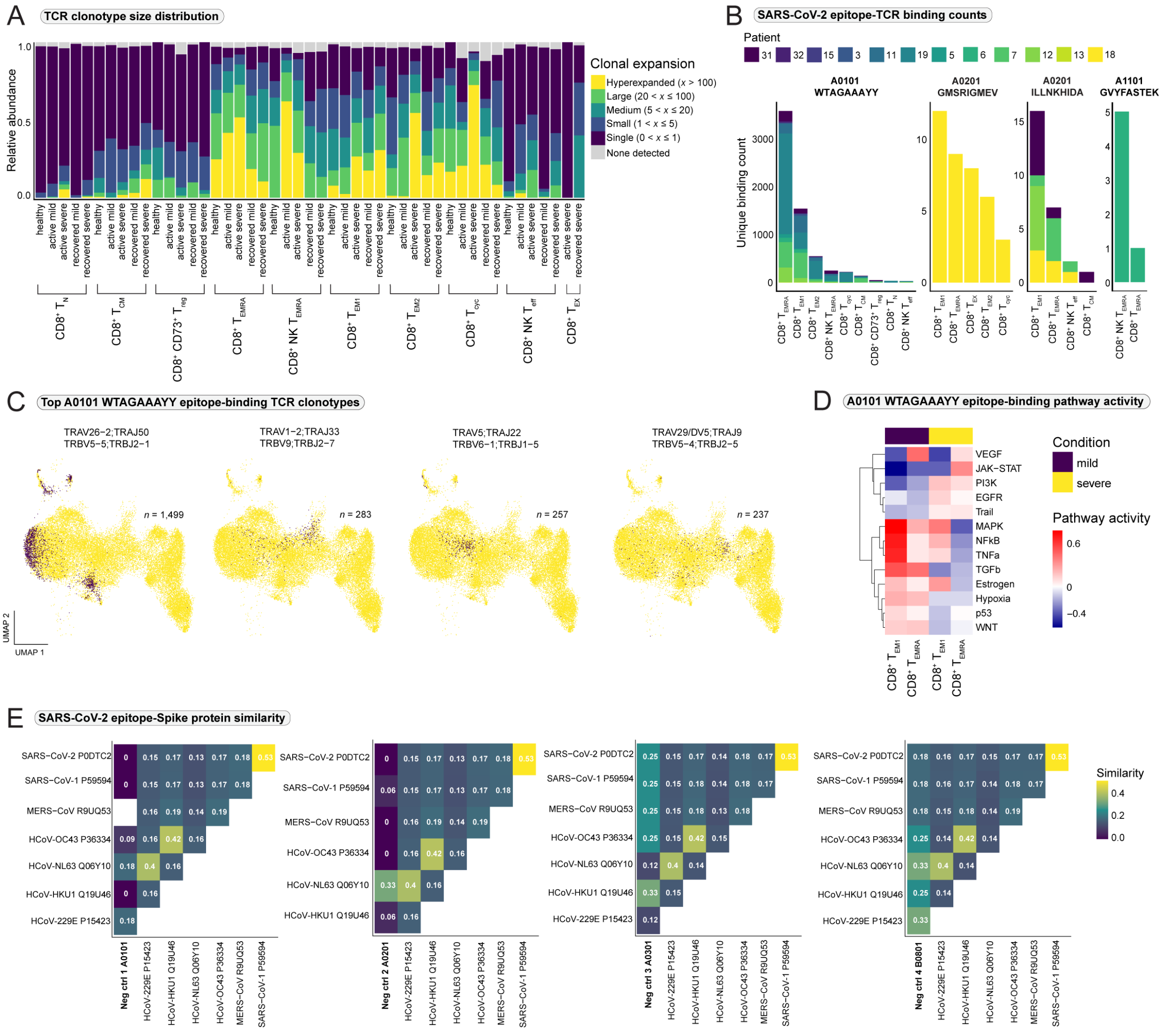


**Supplemental Fig. 5. Additional information on clonotype and epitope-binding analysis, Related to Fig. 4 and 6. (A)** Relative abundance of clonotype expansion groupings within the CD8$^{+}$ T cell populations per condition. **(B)** Binding counts for the four uniquely recognized SARS-CoV-2-derived epitopes per CD8$^{+}$ T cell subtype and patient. **(C)** Top WTAGAAAYY epitope-binding T cell receptor (TCR) clonotypes projected onto the integrated UMAP. **(D)** Estimated pathway activity (PROGENy) in WTAGAAAYY epitope-binding cells. **(E)** Protein similarity between the negative control MHC I Dextramer reagents used in the study (*n* = 4) and Spike proteins from SARS-CoV-2 and other known human coronaviruses.

**Supplemental Table 1. MHC class I Dextramer reagent panel, Related to Fig. 1 and Supplemental Fig. 1**

| HLA allele | HLA supertype | Peptide | DNA barcode | Peptide source |
|---|---|---|---|---|
| A*0101 | A01 | LTDEMIAQY[a] | GGCGTACTTGCGGGCTTC | S Protein |
| A*0101 | A01 | WTAGAAAYY[a] | GTAGACCTCACGAGTAGA | S Protein |
| A*0201 | A02 | TLACFVLAAV[a] | TATATGTGGGACCGAGCT | M Protein |
| A*0201 | A02 | GMSRIGMEV[a] | GCAGGGTGAAGCTAGGTG | N Protein |
| A*0201 | A02 | LLLDRLNQL[a] | GTTCTGCTATATGGTCAC | N Protein |
| A*0201 | A02 | ILLNKHIDA[a] | AGTTGACAAGAGCCGTAC | N Protein |
| A*0201 | A02 | RLNEVAKNL | CGTAACTTGTTGTGAGTC | S Protein |
| A*0201 | A02 | YLQPRTFLL | GACACACTGTGGATGACT | S Protein |
| A*0201 | A02 | VLNDILSRL[a] | AACCAAACTTGCTGATAG | S Protein |
| A*0201 | A02 | NLNESLIDL[a] | GGAAATGACATCTCCGAG | S Protein |
| A*0201 | A02 | FIAGLIAIV[a] | ATTTAGCGGCCGACCGCT | S Protein |
| A*0301 | A03 | RIAGHHLGR | TTTCATCGGGATATGTAA | M Protein |
| A*0301 | A03 | LLNKHIDAYK | ATGAGCGGAGGTAAACGT | N Protein |
| A*0301 | A03 | KTFPPTEPK | GATTACGTGAGAATAACG | N Protein |
| A*0301 | A03 | KSAAEASKK | GACAATTGTGCGCGAGAT | N Protein |
| A*0301 | A03 | RLFRKSNLK | TCGCATCGACAGTCTAGT | S Protein |
| A*0301 | A03 | TLKSFTVEK[a] | TGGGCAAACCAGCCTAAC | S Protein |
| A*0301 | A03 | QIYKTPPIK[a] | AGTGTTCGTAGGGTTGTT | S Protein |
| A*0301 | A03 | MTSCCSCLK | CCCACTGACCTTTCGATG | S Protein |
| A*0301 | A03 | VTYVPAQEK | TGGTGCAAGAGTGTGAGT | S Protein |
| A*1101 | A03 | LSYFIASFR[a] | TGGGAGGTGATGTCATGC | M Protein |
| A*1101 | A03 | KTFPPTEPK | GCCATACACATCTAGGCG | N Protein |
| A*1101 | A03 | KSAAEASKK | CCTTATCTAAAGCGAGTC | N Protein |
| A*1101 | A03 | NSASFSTFK | TGACTGGGCTGGATATGG | S Protein |
| A*1101 | A03 | MTSCCSCLK[a] | TTTACCTGAGTGATATCG | S Protein |
| A*1101 | A03 | VTYVPAQEK | GCTCTCTTTGCCCTGAGG | S Protein |
| A*1101 | A03 | GVYFASTEK[a] | GATCTGCGCATTACGTCA | S Protein |
| A*1101 | A03 | TLKSFTVEK[a] | TTTCTATGTATACAAGAC | S Protein |
| A*1101 | A03 | GVYYHKNNK | GTCCCTGTCACATAGTTA | S Protein |
| A*1101 | A03 | RLFRKSNLK | AAAGAGCTAGCTCAACCT | S Protein |
| A*0101 | A01 | STEGGGLAY[a] | ATGCAGGACTATGTCTTG | Negative control |
| A*0201 | A02 | ALIAPVHAV[a] | CCCTGACGCACCTTATGA | Negative control |
| A*0301 | A03 | GLFGAGAFK[a] | GGCGGGTTCTCCAGCGGA | Negative control |
| B*0801 | B08 | AAKGRGAAL[a] | TGTGACTGCCTCGTAAAC | Negative control |
| A*0101 | A01 | VTEHDTLLY[a] | CTTTGTAGTACGACGTCA | CMV pp50 |
| A*0201 | A02 | NLVPMVATV[a] | GATGCTAGTGCACGTCTA | CMV pp65 |
| A*0301 | A03 | KLGGALQAK[a] | CTTTGGTGGAGGCGCAGT | CMV IE-1 |
| A*1101 | A03 | ATVQGQNLK[a] | GTGGCGAATTTGACTCGT | CMV pp65 |

[a] MHC class I Dextramer reagents selected for the single-cell experiment

### Supplemental Table 2. Clinical data, Related to Fig. 1 and Methods

| patient | sex | age | ICU | ventilation | days since first positive PCR | group | follow up |
|---|---|---|---|---|---|---|---|
| P01[a] | female | 78 | no | no | 5 | active mild | recovered |
| P02[a] | female | 77 | no | no | 7 | active mild | recovered |
| P03[a] | male | 64 | yes | yes | 12 | active severe | recovered |
| P04 | male | 71 | no | no | 2 | active mild | recovered |
| P05[a] | female | 27 | no | no | 50 | recovered mild | recovered |
| P06[a] | male | 28 | no | no | 58 | recovered mild | recovered |
| P07[a] | female | 25 | no | no | 57 | recovered mild | recovered |
| P10 | male | 53 | yes | yes | 53 | recovered severe | recovered |
| P11[a] | male | 56 | yes | yes | 57 | active severe | recovered |
| P12[a] | female | 67 | yes | yes | 55 | recovered severe | deceased |
| P13[a] | female | 62 | yes | yes | 49 | recovered severe | deceased |
| P14 | male | 55 | yes | yes | na | active severe | deceased |
| P15[a] | female | 75 | no | no | 7 | active mild | recovered |
| P16 | male | 67 | yes | yes | 52 | recovered severe | recovered |
| P17 | female | 56 | yes | yes | 44 | recovered severe | recovered |
| P18[a] | male | 62 | yes | yes | 96 | recovered severe | deceased |
| P19[a] | male | 82 | yes | yes | 11 | active severe | recovered |
| P20 | male | 67 | yes | yes | na | recovered severe | recovered |
| P21 | female | 77 | yes | yes | 25 | recovered severe | recovered |
| P22 | male | 48 | yes | yes | 45 | recovered severe | deceased |
| P23 | male | 68 | yes | yes | na | recovered severe | deceased |
| P24 | male | 59 | yes | yes | 5 | recovered severe | recovered |
| P26 | female | 35 | - | - | - | healthy | healthy |
| P27 | female | 51 | - | - | - | healthy | healthy |
| P29[a] | female | 52 | - | - | - | healthy | healthy |
| P30 | female | 40 | - | - | - | healthy | healthy |
| P31[a] | male | 38 | - | - | - | healthy | healthy |
| P32[a] | male | 79 | - | - | - | healthy | healthy |

[a] Patients, who were selected for the single-cell experiment. ICU = intensive care unit, na = not available

**Supplemental Table 3. Patient HLA alleles and related HLA supertypes, Related to Fig. 1 and Supplemental Fig. 1**

| | HLA-A allele 1 | HLA-A supertype 1 | HLA-A allele 2 | HLA-A supertype 2 | HLA-B allele 1 | HLA-B supertype 1 | HLA-B allele 2 | HLA-B supertype 2 |
|---|---|---|---|---|---|---|---|---|
| P01 | 01:01P | A01 | 03:01P | A03 | 07:02P | B07 | 08:01P | B08 |
| P02 | 02:01P | A02 | - | na | 40:01P | B44 | 44:03:01 | B44 |
| P03 | 02:01P | A02 | 26:01P | A01 | 35:08P | B07 | 51:01P | B07 |
| P04 | 02:01:01G | A02 | - | na | 07:02:01G | B07 | 51:01:01G | B07 |
| P05 | 02:01P | A02 | - | na | 40:01P | B44 | 48:01P | B27 |
| P06 | 29:02P | A01A24 | 31:01P | A03 | 44:03:01 | B44 | 58:01P | B58 |
| P07 | 02:01P | A02 | - | na | 18:01P | B44 | 44:02P | B44 |
| P10 | 03:01P | A03 | 26:01P | A01 | 07:02P | B07 | 14:01:01 | B27 |
| P11 | 03:01P | A03 | 68:01P | A03 | 13:02P | unclassified | 44:02P | B44 |
| P12 | 03:01P | A03 | 68:01P | A03 | 07:02P | B07 | 40:01P | B44 |
| P13 | 01:01P | A01 | 30:02P | A03 | 07:02P | B07 | 49:01P | unclassified |
| P14 | 02:01:01G | A02 | 11:01:01G | A03 | 18:01:01G | B44 | 37:01:01G | B44 |
| P15 | 02:01P | A02 | - | na | 44:02P | B44 | - | na |
| P16 | 01:01P | A01 | 66:01:01 | A03 | 35:01P | B07 | 35:03P | B07 |
| P17 | 03:01P | A03 | 68:02P | A02 | 07:02P | B07 | 51:01P | B07 |
| P18 | 11:01P | A03 | 25:01P | A01 | 44:02P | B44 | 58:01P | B58 |
| P19 | 02:01:01G | A02 | 03:01:01G | A03 | 07:02:01G | B07 | 27:05:02G | B27 |
| P20 | 03:01P | A03 | 26:01P | A01 | 38:01:01 | B27 | 51:01P | B07 |
| P21 | 01:01P | A01 | 03:01P | A03 | 08:01P | B08 | 51:01P | B07 |
| P22 | 03:01P | A03 | 24:02P | A24 | 15:24:01 | B62 | 57:01P | B58 |
| P23 | 01:01P | A01 | - | na | 07:02P | B07 | 57:01P | B58 |
| P24 | 02:01P | A02 | 31:01P | A03 | 39:01P | B27 | 56:01P | B07 |
| P26 | 24:02P | A24 | 29:02P | A01A24 | 15:07P | B62 | 58:01P | B58 |
| P27 | 03:01P | A03 | - | na | 07:02P | B07 | 35:01P | B07 |
| P29 | 02:05:01:01 | A02 | 31:01P | A03 | 51:01P | B07 | 58:01P | B58 |
| P30 | 03:01P | A03 | 24:02P | A24 | 18:01P | B44 | 35:01P | B07 |
| P31 | 02:01P | A02 | 33:01:01 | A03 | 07:02P | B07 | 27:05P | B27 |
| P32 | 01:01P | A01 | 02:01P | A02 | 14:01:01 | B27 | 58:01P | B58 |

P = HLA alleles with an identical protein sequence in their peptide-binding domain, G = HLA alleles with an identical nucleotide sequence encoding for their peptide-binding domain, unclassified = the respective HLA allele has not been assigned to an HLA supertype, na = not available

**Supplemental Table 4. Marker genes and antibody-derived tags (adt) used for annotation of $CD8^+$ T cell subtypes, Related to Fig. 1 and Methods.**

**Supplemental Table 5. Overview of all differentially expressed genes, Related to Fig. 2, 3, and 6.**
***Sheet 1***: Differentially expressed genes between active severe and active mild COVID-19. (*FindMarkers*, *Seurat*). Positive $\log_2$ fold changes indicate upregulation in active severe COVID-19.
***Sheet 2***: Differentially expressed genes between active mild COVID-19 and healthy controls (*FindMarkers*, *Seurat*). Positive $\log_2$ fold changes indicate upregulation in active mild COVID-19.
***Sheet 3***: Differentially expressed genes between active severe COVID-19 and healthy controls (*FindMarkers*, *Seurat*). Positive $\log_2$ fold changes indicate upregulation in active severe COVID-19.
***Sheet 4***: Differentially expressed genes between recovered severe and recovered mild COVID-19 (*FindMarkers*, *Seurat*). Positive $\log_2$ fold changes indicate upregulation in recovered severe COVID-19.
***Sheet 5***: Differential gene expression between the progenitor and differentiated cell populations (*startVsEndTest*, *tradeSeq*). Positive $\log_2$ fold changes indicate upregulation in the differentiated cell populations.
***Sheet 6***: Differential expression at the bifurcation point of the SLEC and MPEC trajectories (*earlyDETest*, *tradeSeq*). Positive $\log_2$ fold changes indicate upregulation in the MPEC lineage.
***Sheet 7***: Differential gene expression between the end stages of the lineages (*diffEndTest*, *tradeSeq*). Positive $\log_2$ fold changes indicate upregulation in the SLEC lineage.
*Sheet 8*: Differential gene expression between active severe and mild COVID-19 across pseudotime (*conditionTest*, *tradeSeq*).
***Sheet 9***: Differentially expressed genes between WTAGAAAYY epitope-binding $CD8^+$ T cells in severe and mild COVID-19 (*FindMarkers*, *Seurat*). Positive $\log_2$ fold changes indicate upregulation in severe COVID-19.

**Supplemental Table 6. Absolute counts for epitope-binding cells passing filter, Related to Fig. 4 and Supplemental Fig. 5**

| HLA allele and epitope | number of binding cells | number of uniquely binding cells |
|---|---|---|
| A*0101 WTAGAAAYY | 7300 | 6431 |
| A*0201 GMSRIGMEV | 365 | 38 |
| A*0201 LLLDRLNQL | 9 | 0 |
| A*0201 ILLNKHIDA | 812 | 26 |
| A*0201 FIAGLIAIV | 8 | 0 |
| A*1101 LSYFIASFR | 322 | 0 |
| A*1101 GVYFASTEK | 106 | 6 |
| A*1101 TLKSFTVEK | 11 | 0 |
| total binding cells | 7684 | 6501 |

### Supplemental Table 7. Primer sequences for bulk sequencing, Related to Methods

| PCR step | primer | oligonucleotide sequence |
|---|---|---|
| 1. PCR | forward primer | CTCTTTCCCTACACGACGCTCTTCCGATCTGAAGTTCCAGCCAGCGTC |
| | reverse primer | CTGGAGTTCAGACGTGTGCTCTTCCGATCTCTGTGACTATGTGAGGCTTTC |
| 2. PCR | common primer | AATGATACGGCGACCACCGAGATCTACACTCTTTCCCTACACGACGCTCTTCCGATCT |
| | barcoded primer 1 | CAAGCAGAAGACGGCATACGAGATAACGTGATGTGACTGGAGTTCAGACGTGTGCTCTTCCGATCT |
| | barcoded primer 2 | CAAGCAGAAGACGGCATACGAGATAAACATCGGTGACTGGAGTTCAGACGTGTGCTCTTCCGATCT |
| | barcoded primer 3 | CAAGCAGAAGACGGCATACGAGATATGCCTAAGTGACTGGAGTTCAGACGTGTGCTCTTCCGATCT |
| | barcoded primer 4 | CAAGCAGAAGACGGCATACGAGATAGTGGTCAGTGACTGGAGTTCAGACGTGTGCTCTTCCGATCT |
| | barcoded primer 5 | CAAGCAGAAGACGGCATACGAGATCAGATCTGGTGACTGGAGTTCAGACGTGTGCTCTTCCGATCT |
| | barcoded primer 6 | CAAGCAGAAGACGGCATACGAGATCATCAAGTGTGACTGGAGTTCAGACGTGTGCTCTTCCGATCT |
| | barcoded primer 7 | CAAGCAGAAGACGGCATACGAGATCGCTGATCGTGACTGGAGTTCAGACGTGTGCTCTTCCGATCT |
| | barcoded primer 8 | CAAGCAGAAGACGGCATACGAGATCTGTAGCCGTGACTGGAGTTCAGACGTGTGCTCTTCCGATCT |
| | barcoded primer 9 | CAAGCAGAAGACGGCATACGAGATGACTAGTAGTGACTGGAGTTCAGACGTGTGCTCTTCCGATCT |
| | barcoded primer 10 | CAAGCAGAAGACGGCATACGAGATGAATCTGAGTGACTGGAGTTCAGACGTGTGCTCTTCCGATCT |
| | barcoded primer 11 | CAAGCAGAAGACGGCATACGAGATGAGCTGAAGTGACTGGAGTTCAGACGTGTGCTCTTCCGATCT |
| | barcoded primer 12 | CAAGCAGAAGACGGCATACGAGATGATAGACAGTGACTGGAGTTCAGACGTGTGCTCTTCCGATCT |
| | barcoded primer 13 | CAAGCAGAAGACGGCATACGAGATTAGGATGAGTGACTGGAGTTCAGACGTGTGCTCTTCCGATCT |
| | barcoded primer 14 | CAAGCAGAAGACGGCATACGAGATTATCAGCAGTGACTGGAGTTCAGACGTGTGCTCTTCCGATCT |
| | barcoded primer 15 | CAAGCAGAAGACGGCATACGAGATTCCGTCTAGTGACTGGAGTTCAGACGTGTGCTCTTCCGATCT |
| | barcoded primer 16 | CAAGCAGAAGACGGCATACGAGATTCTTCACAGTGACTGGAGTTCAGACGTGTGCTCTTCCGATCT |
| | barcoded primer 17 | CAAGCAGAAGACGGCATACGAGATACCACTGTGTGACTGGAGTTCAGACGTGTGCTCTTCCGATCT |
| | barcoded primer 18 | CAAGCAGAAGACGGCATACGAGATACATTGGCGTGACTGGAGTTCAGACGTGTGCTCTTCCGATCT |
| | barcoded primer 19 | CAAGCAGAAGACGGCATACGAGATACAAGCTAGTGACTGGAGTTCAGACGTGTGCTCTTCCGATCT |
| | barcoded primer 20 | CAAGCAGAAGACGGCATACGAGATAGTACAAGGTGACTGGAGTTCAGACGTGTGCTCTTCCGATCT |
| | barcoded primer 21 | CAAGCAGAAGACGGCATACGAGATCAACCACAGTGACTGGAGTTCAGACGTGTGCTCTTCCGATCT |
| | Barcodes primer 22 | CAAGCAGAAGACGGCATACGAGATCAATGGAAGTGACTGGAGTTCAGACGTGTGCTCTTCCGATCT |
| | barcoded primer 23 | CAAGCAGAAGACGGCATACGAGATCACTTCGAGTGACTGGAGTTCAGACGTGTGCTCTTCCGATCT |
| | barcoded primer 24 | CAAGCAGAAGACGGCATACGAGATCAGCGTTAGTGACTGGAGTTCAGACGTGTGCTCTTCCGATCT |

Blue = overhang sequences that function as a handle sequence for primers used in the second PCR-step, green = individual index-sequences
